# Actin polymerization drives lumen formation in a human epiblast model

**DOI:** 10.1101/2023.04.20.537711

**Authors:** Dhiraj Indana, Andrei Zakharov, Youngbin Lim, Alexander R. Dunn, Nidhi Bhutani, Vivek B. Shenoy, Ovijit Chaudhuri

## Abstract

Lumens or fluid-filled cavities are a ubiquitous feature of mammals and are often evolutionarily linked to the origin of body-plan complexity. Post-implantation, the pluripotent epiblast in a human embryo forms a central lumen, paving the way for gastrulation. While osmotic pressure gradients drive lumen formation in many developmental contexts, mechanisms of human epiblast lumenogenesis are unknown. Here, we study lumenogenesis in a pluripotent-stem-cell-based model of the epiblast using engineered hydrogels that model the confinement faced by the epiblast in the blastocyst. Actin polymerization into a dense mesh-like network at the apical surface generates forces to drive early lumen expansion, as leaky junctions prevent osmotic pressure gradients. Theoretical modeling reveals that apical actin polymerization into a stiff network drives lumen opening, but predicts that a switch to pressure driven lumen growth at larger lumen sizes is required to avoid buckling of the cell layer. Consistent with this prediction, once the lumen reaches a radius of around 12 μm, tight junctions mature, and osmotic pressure gradients develop to drive further lumen growth. Human epiblasts show a transcriptional signature of actin polymerization during early lumenogenesis. Thus, actin polymerization drives lumen opening in the human epiblast, and may serve as a general mechanism of lumenogenesis.

## Introduction

During human embryonic development, the fertilized egg undergoes multiple rounds of cell division and differentiation to form the pluripotent epiblast at the blastocyst stage, which ultimately gives rise to all tissues in the fetus (Rossant and Tam, 2017). Embryo development from the pluripotent epiblast commences upon implantation of the blastocyst into the uterine wall, following which pluripotent stem cells self-organize to form a roughly spherical structure containing a fluid-filled lumen (Shahbazi and Zernicka-Goetz, 2018). Proper formation of the epiblast lumen is critical for establishing morphogen gradients that drive subsequent embryonic development (Zhang et al., 2019). While the physical mechanism of epiblast lumenogenesis is unknown, established mechanisms of de novo lumenogenesis in other model systems involve apoptosis and osmotic pressure gradients (Blasky et al., 2015; Sigurbjörnsdóttir et al., 2014). Apoptosis drives lumenogenesis in certain mammary epithelial models where cells at the center of a cluster die, resulting in a hollow cavity (Debnath et al., 2002). Osmotic pressure gradients drive lumen growth in the mouse blastocyst (Chan et al., 2019; Dumortier et al., 2019; Ruiz-Herrero et al., 2017), MDCK (Madin-Darby canine kidney) cells (Latorre et al., 2018), bile canaliculi (Dasgupta et al., 2018), and zebrafish inner ear (Mosaliganti et al., 2019). In each of these cases, apico-basally polarized cells with tight junctions, pump osmolytes into the lumen which builds osmotic pressure and drives water into the lumen, expanding its volume (Chugh et al., 2022; Torres-Sánchez et al., 2021). While pressure has been shown to drive lumenogenesis in the mouse blastocyst (Chan et al., 2019; Dumortier et al., 2019; Ruiz-Herrero et al., 2017), mechanisms of lumenogenesis in other early embryonic lumens such as the epiblast cavity are much less understood (Carleton et al., 2022; Kim and Bedzhov, 2022). Recently, the study of polarity (Wang et al., 2021) and pluripotency (Shahbazi et al., 2017) dynamics necessary for epiblast lumenogenesis have provided key insights into the cellular processes involved, but the physics driving lumenogenesis in the human epiblast remains unclear.

Human induced pluripotent stem cell (hiPSC) models of the embryo reproduce key aspects of development and serve as excellent tools to uncover mechanisms orchestrating human embryogenesis (Bao et al., 2022; Fu et al., 2021), since human embryos cannot be studied directly due to ethical concerns. We have previously shown that hiPSCs form lumen-containing structures that morphologically and phenotypically model the human epiblast in a highly reproducible manner, when cultured in 3D in engineered hydrogels which model the confinement experienced by the epiblast in vivo due to blastocyst cavity pressure (Chan et al., 2019) and extraembryonic cells (Indana et al., 2021). In this epiblast model (hereafter referred to as synthetic epiblast), we now dissect the mechanisms regulating lumen formation. Our experiments and simulations reveal a previously undescribed mechanism of lumen formation mediated by apical actin polymerization that drives early lumen opening till a critical lumen size of ~12 μm radius, followed by a transition to osmotic pressure gradient driven lumen growth in lumens larger than 12 μm radius.

## Results

### hiPSCs form synthetic epiblasts in 3D hydrogels

We formed synthetic epiblasts by culturing single hiPSCs in 3D viscoelastic alginate hydrogels. In the presence of optimal biophysical cues of hydrogel stiffness, viscoelasticity, and cell-adhesion ligand density, hiPSCs self-organize into lumen-containing structures that are reminiscent of the human epiblast (Indana et al., 2021). While the initial elastic modulus of the hydrogels is 20 kPa, these gels exhibit fast stress relaxation, with a stress relaxation half time of ~70 s (Fig. S1A). Thus, the relaxation modulus over ~30 mins is expected to be ~1 kPa, which is on the same order of magnitude as that experienced by the epiblast cells in the blastocyst (Chan et al., 2019) (Fig. S1A). In these hydrogels, hiPSCs proliferate and begin to form lumens around day 3 of culture, creating 3D monolayered cellular structures with a central, roughly spherical lumen (Fig. 1, A and B). Similar to human epiblasts (Deglincerti et al., 2016; Shahbazi et al., 2016; Shahbazi et al., 2017), these structures polarized along the apical-basal axis in response to matrix signaling (Indana et al., 2021) and maintained expression of core pluripotency proteins such as Oct4, Sox2 and Nanog, as well as primed pluripotency factors such as Otx2, over 7 days of culture (Fig. 1C, Fig. S1, B and C, and Video S1). hiPSC clusters also mirrored the morphological features of human epiblasts. The numbers of cells in hiPSC clusters on different days of culture were akin to human epiblasts, with day 3 and day 7 of in vitro culture corresponding to 7 to 8 days post fertilization (d.p.f.) and 11 to 12 d.p.f in human embryos respectively, suggesting similar proliferation dynamics (Fig. 1D). Further, lumen formation began in clusters of ~4-5 cells in both and, lumen volumes and growth rates of hiPSC clusters were close to those of human epiblasts (Fig. 1E). Nuclear morphology metrics such as area and perimeter of hiPSC clusters were also similar to those of human epiblasts (Fig. S1D). Overall, hiPSCs in optimal 3D alginate hydrogels maintained pluripotency, polarized along the apical-basal axis, and showed lumenal and nuclear morphological similarities to human epiblasts. Therefore, these hiPSC-based structures serve as synthetic epiblasts, allowing study of mechanisms driving epiblast lumenogenesis.

**Figure 1.**
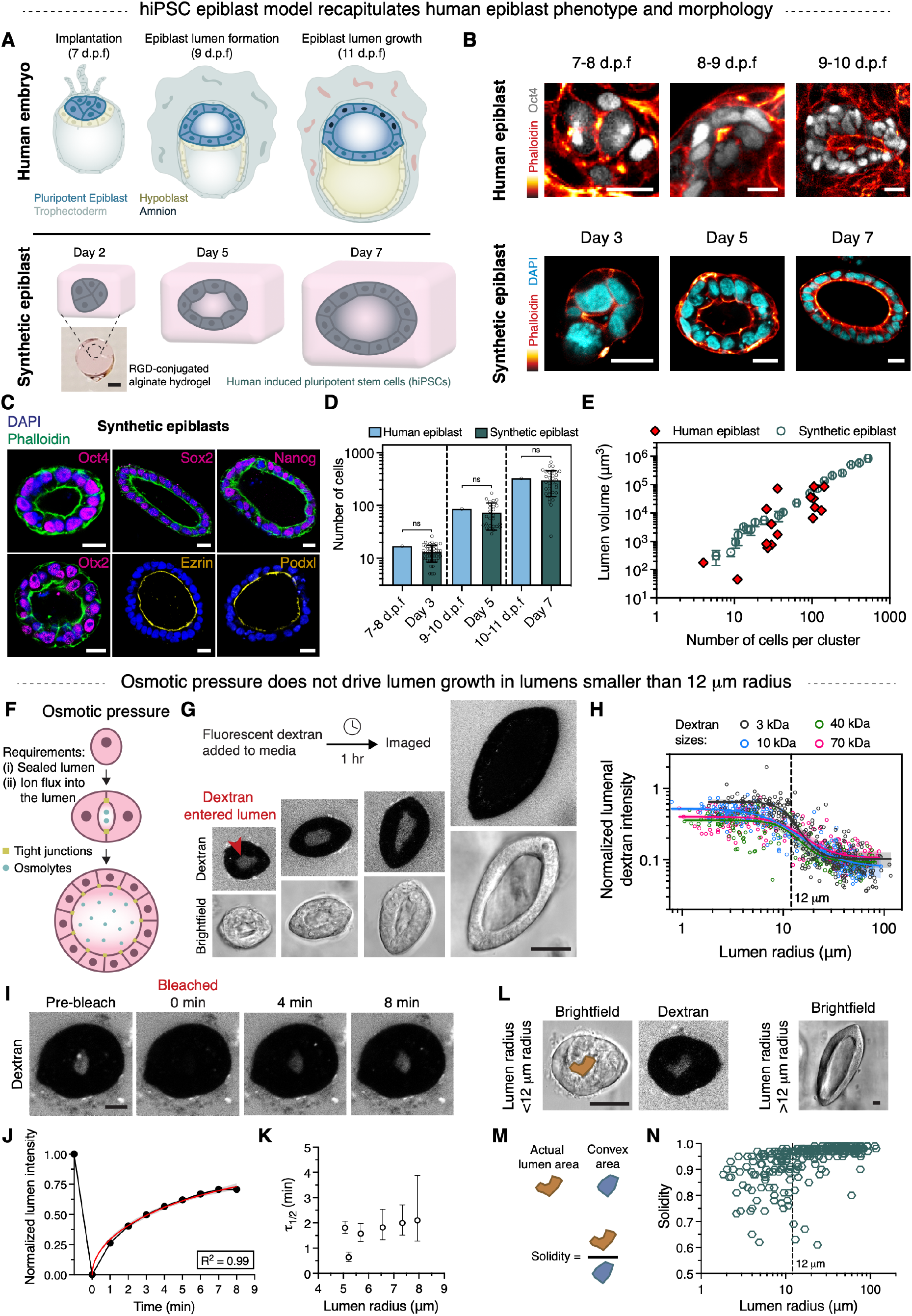
Synthetic epiblasts model the human epiblast and osmotic pressure does not drive lumenogenesis in synthetic epiblasts. **(A)** Schematic of peri-implantation human embryos and synthetic epiblasts. **(B)** Immunostains of human and synthetic epiblasts. Human epiblast images were generated in and modified with permission from ref. (Shahbazi et al., 2016). **(C)** Immunostains of Oct4, Sox2, Nanog (core pluripotency), Otx2 (primed pluripotency), Ezrin and Podocalyxin (apical polarity) in synthetic epiblasts. **(D)** Quantification of cell numbers in human (Shahbazi et al., 2016) and synthetic epiblasts (mean ± s.d.; ns: not significant p > 0.05, one-way ANOVA; *n* = 43 (day 3), 26 (day 5), 32 (day 7) synthetic epiblasts). Data for human epiblasts was generated in ref (Shahbazi et al., 2016). **(E)** Correlation between lumen volume and cell numbers in human (Deglincerti et al., 2016; Molè et al., 2021; Shahbazi et al., 2016; Shahbazi et al., 2017; Simunovic et al., 2019) and synthetic epiblasts (mean ± s.d. for synthetic epiblasts; *n* = 16 (human), 101 (synthetic)). **(F)** Schematic of osmotic pressure driven lumenogenesis. **(G)** Tight junction permeability assay. Fluorescence images of cell-impermeant dextran. **(H)** Quantification of dextran intensity inside lumen. Lines indicate sigmoidal fits and 95% CI. (*n*, R^2^) = 3 kDa: (290, 0.71), 10 kDa: (229, 0.81), 40 kDa: (213, 0.69), 70 kDa: (228, 0.69). **(I)** Fluorescence recovery after photobleaching (FRAP) of lumenal dextran. **(J)** Normalized lumenal dextran intensity quantification for cluster shown in (I). Line (red) indicates least squares fit based on a FRAP model (Soumpasis, 1983) and 95% CI. **(K)** Estimation of fluorescence recovery half-time (τ_1/2_) (mean ± 95% CI). **(L-N)** Quantification of lumen solidity. (*n* = 290 for plot (N)). Scale bars: 2 mm (A), 20 μm (B, C), 50 μm (G), 10 μm (I), 25 μm (L).

### Osmotic pressure gradients and apoptosis do not drive lumen opening

We sought to understand the mechanisms underlying lumenogenesis in synthetic epiblasts as a model of human epiblasts. Guided by previous studies of lumenogenesis (Blasky et al., 2015; Sigurbjörnsdóttir et al., 2014), we first investigated known mechanisms of de novo lumenogenesis including apoptosis and osmotic pressure gradients. During lumen formation and growth in synthetic epiblasts, no apoptotic cells were detected (Fig. S1, E to G), demonstrating that apoptosis does not drive lumenogenesis in synthetic epiblasts, consistent with previous studies (Bedzhov and Zernicka-Goetz, 2014).

With apoptosis eliminated, we next studied the role of osmotic pressure gradients in driving lumenogenesis. To build osmotic pressure in the lumen, two requirements need to be met: (i) ion flux into intercellular space or lumen at the apical surface (Torres-Sánchez et al., 2021), and (ii) formation of tight junctions to prevent osmolytes from leaking (Shen et al., 2011) (Fig. 1F). These requirements allow osmotic pressure to build up, which draws water into the lumen, generating force necessary for lumen growth. To test if hiPSCs formed tight junctions, cell-impermeable fluorescent dextran was added to the culture media. If functional tight junctions were present, dextran would be expected to be excluded from the lumen. Strikingly, dextran entered lumens smaller than ~12 μm in radius, indicating that hiPSCs do not form functional tight junctions during early stages of lumenogenesis when the lumen size is below ~12 μm radius (Fig. 1, G and H, and Fig. S2, A to E). Dextran also localized to the intercellular spaces in junctions between cells and was excluded from cells themselves suggesting that dextran entered lumens through diffusion along the intercellular spaces and not via other mechanisms such as transcytosis (Fig. S2, F and G). As tight junction marker ZO-1 localized to the cell-cell boundary of smaller lumens as well (Fig. S2H, and Video S1), it was plausible that tight junction formation was gradual with a complete seal forming at a lumen size of ~12 μm radius. But this was not the case. Large macromolecular dextran, with a diameter (~12 nm; 70 kDa dextran) comparable to intercellular space due to adherens junctions (~20 nm) (Farquhar and Palade 1965; Otani et al., 2019), entered lumens smaller than ~12 μm in radius but was excluded from lumens larger than this size, highlighting the complete lack of functional tight junctions in synthetic epiblasts with smaller lumens (Fig. 1H, and Fig. S2, D and E).

As functional tight junctions were absent in synthetic epiblasts with smaller lumens (radius < 12 μm; hereafter referred to as smaller epiblasts), any ions pumped into these lumens are expected to leak along the intercellular spaces, preventing large pressures from building up. To confirm that this was the case, diffusion dynamics in smaller lumens were measured by observing fluorescence recovery after photobleaching (FRAP) of dextran. The fluorescence signals recovered ~2 min after photobleaching, suggesting that dextran can freely diffuse along the intercellular spaces (Fig. 1, I to K, and Video S2). Taken together, these results reveal a lumen size-dependent initiation of tight junction formation, with lumens below ~12 μm in radius being leaky.

To further assess the role of osmotic pressure gradients in driving lumenogenesis, lumen shapes were examined. Lumen shapes are expected to be convex or bent outward if pressure was the sole driver of epiblast lumenogenesis, whereas irregularly shaped lumens that are bent inwards suggest that osmotic pressure gradients are not a dominant driver of lumen growth (Vasquez et al., 2021). Lumen shapes were highly irregular for smaller lumens but transitioned to a more bulged, convex shape in larger lumens (Fig. 1, L to N). Thus, the irregularly shaped lumens in smaller epiblasts further indicate that osmotic pressure is not a major driver of early lumenogenesis in synthetic epiblasts, whereas the regularly shaped lumens in larger epiblasts indicate that osmotic pressure gradients could drive lumen expansion in larger lumens. Finally, laser ablation through an entire cell in smaller epiblasts did not cause any drastic change in cell or lumen size or shape indicating that smaller lumens are not pressurized (Fig. S2I). Taken together, the lack of tight junctions, free diffusion out of the intralumenal space, and lumen shapes together indicate that initial expansion of the lumen is not driven by osmotic pressure gradients and emphasize the existence of a previously undescribed pressure-independent mechanism of lumenogenesis.

### Early lumenogenesis is associated with force generation and formation of an apical actin mesh

As epiblast lumenogenesis mechanisms are required to produce forces necessary to overcome resistance from their environment – the surrounding hydrogel in case of the synthetic epiblast versus extraembryonic cells and blastocyst cavity pressure (Chan et al., 2019) in case of the human epiblast – we next examined force generation associated with lumenogenesis in order to gain insight into the underlying mechanisms driving early lumen expansion. Synthetic epiblasts of all sizes generated local matrix deformations on the order of tens of micrometers over 18 hr (Fig. 2, A and B, and Videos S3 and S4). As the hydrogels are viscoelastic and viscoplastic, with stresses relaxed on a timescale of minutes and the material undergoing permanent deformation, the magnitude of forces required for the measured matrix deformations depends on the timescale and dynamics of force application. Nonetheless, as some force generation is necessary, we next probed different cellular force generating machineries to uncover the pressure-independent mechanism responsible for epiblast lumenogenesis.

**Figure 2.**
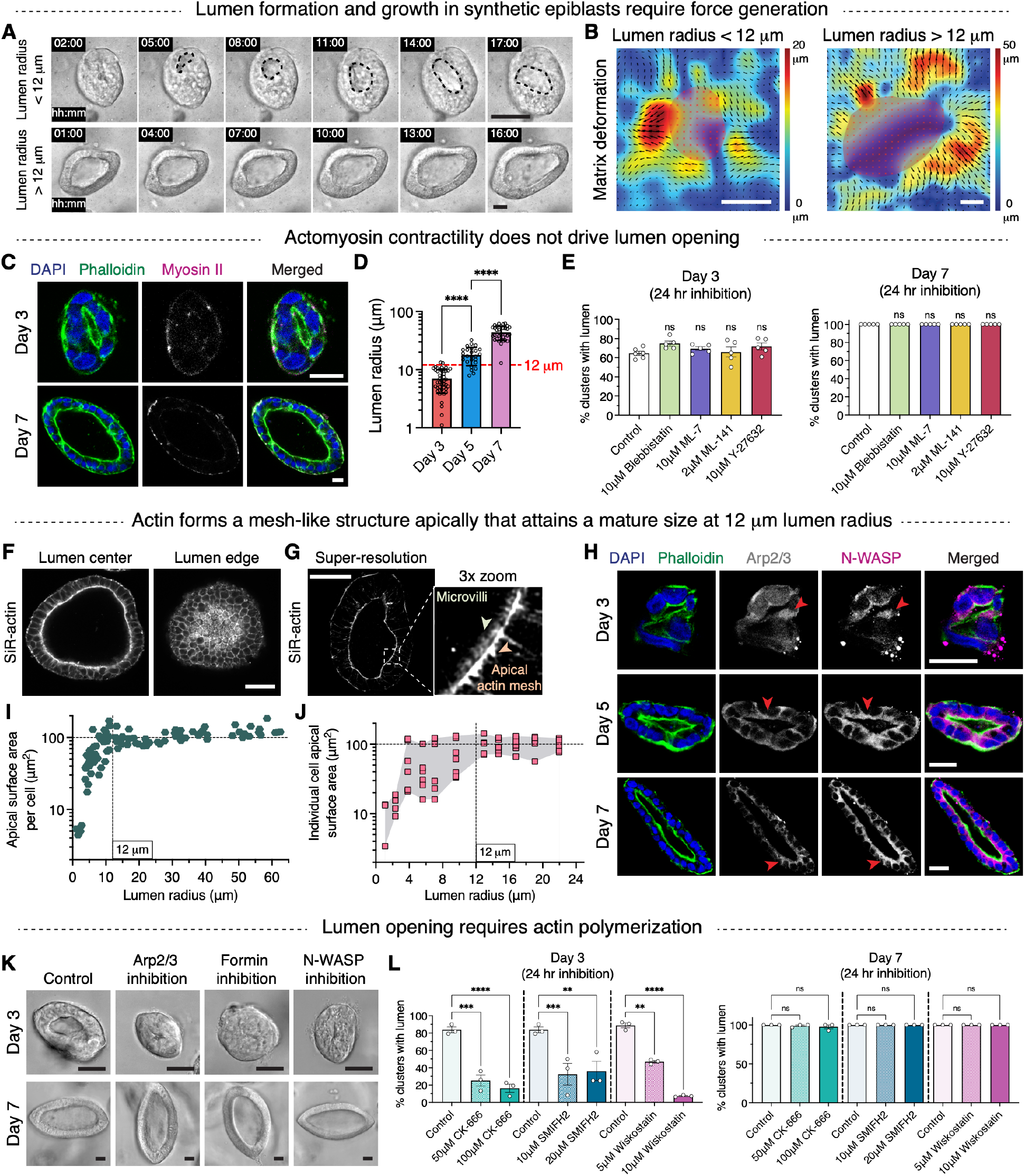
Early lumenogenesis is associated with force generation and apical actin mesh formation. **(A)** Timelapse (hh:mm) images of lumen opening and growth. Black outlines indicate luminal surface. **(B)** Matrix deformations generated during lumen growth for clusters shown in (A). **(C)** Immunostains of myosin II, phalloidin (actin) and DAPI (nucleus) in synthetic epiblasts. **(D)** Quantification of lumen radius on different days of culture (mean ± s.d.; ****p < 0.0001, one-way ANOVA; *n* = 43 (day 3), 26 (day 5), 32 (day 7)). **(E)** Percent epiblasts with lumen in the presence of different actomyosin contractility inhibitors (mean ± s.e.m.; ns: not significant p > 0.05, one-way ANOVA; *n* = 5). **(F and G)** Representative confocal (F) and super-resolution (G) images of apical actin mesh. **(H)** Immunostains of Arp2/3 and N-WASP in synthetic epiblasts. Red arrowheads highlight apical localization of Arp2/3 and N-WASP. **(I)** Quantification of average apical surface area per cell as a function of lumen size (*n* = 101 synthetic epiblasts). **(J)** Quantification of apical surface area of individual cells in an epiblast. Shaded (gray) region indicates range (max-min) of areas per epiblast (*n* = 11 synthetic epiblasts; ≥ 3 cells per synthetic epiblast). **(K and L)** Representative brightfield images (K) and quantification of percent epiblasts with lumen (L) in the presence of Arp2/3 (CK-666), formin (SMIFH2) and N-WASP (Wiskostatin) inhibitors (mean ± s.e.m.; ****p < 0.0001, ***p < 0.001, **p < 0.01, ns: not significant p > 0.05, one-way ANOVA; *n* = 3). Scale bars: 40 μm (A and B), 20 μm (C), 50 μm (F and G), 20 μm (H), 25 μm (K).

We first examined the role of actomyosin contractility in lumenogenesis, given the well-known function of the actomyosin cytoskeleton network in generating contractile forces. Myosin II was mostly punctate and localized at the basal surface, which could not explain the pattern of forces associated with lumenogenesis (Fig. 2C, and Fig. S3A). Further, inhibition of actomyosin contractility on day 3 of culture or in smaller epiblasts (lumen radius < 12 μm) as well as on day 7 or in larger epiblasts (lumen radius > 12 μm) (Fig. 2D), did not significantly impact lumen formation (Fig. 2E and Fig. S3B). These results demonstrate that actomyosin contractility does not drive lumen formation in synthetic epiblasts.

We next examined actin structures and their potential role in driving lumenogenesis, as actin polymerization in bundled or branched networks produces protrusive forces that drive cellular morphogenesis in a variety of contexts (Fletcher and Mullins, 2010). In synthetic epiblasts, actin was densely localized at the apical surface (Fig. 2F). Super-resolution microscopy using an Airyscan system revealed that apically, actin formed a dense mesh-like structure with microvilli protruding from this mesh (Fig. 2G and Video S5). Further, actin nucleation factor N-WASP and actin branching complex Arp2/3 were enriched at the apical surface, which would be expected to promote the formation of a dendritic actin network (Fig. 2H). As formation of a lumen and apical surface are intertwined, the time evolution of the apical actin mesh formation was quantified. Apical surface area per cell increased in size as lumens grew but reached an equilibrium size of ~100 μm^2^ at a lumen size of ~12 μm radius (Fig. 2I and Fig. S3, C and D). In fact, all cells in larger epiblasts had a similar apical surface area of ~100 μm^2^ while cells in smaller epiblasts had a wide range of apical surface areas at any given timepoint that were close to or less than 100 μm^2^, highlighting cell-cell variations in apical surface formation and therefore actin polymerization rates (Fig. 2J). Distinct lumen growth dynamics were observed for smaller and larger lumens while cell volume and thickness stayed relatively constant (Fig. S3, E to G). Overall, these data show that as lumens form, cells grow their apical surfaces up to an equilibrium value, which is achieved at a lumen size of ~12 μm radius, coinciding with the timing of tight junction formation.

We next tested whether actin polymerization could drive early lumenogenesis using inhibition studies. Dendritic actin network growth is driven by the Arp2/3 complex, which is nucleated via N-WASP, while linear actin polymerization is initiated via formins. Strikingly, inhibition of actin polymerization by any of these proteins – Arp2/3 complex, N-WASP, and formins – strongly reduced lumen formation in smaller epiblasts but had no impact on larger epiblasts (Fig. 2, K and L, and Fig. S3, H and I). Thus, actin polymerization is necessary for lumen formation in smaller epiblasts. Given these observations, we hypothesized that the growth of apical actin in each cell generates force to drive epiblast lumenogenesis in a pressure-independent manner until apical actin growth stalls at a lumen radius of ~12 μm.

### Apical actin polymerization drives lumenogenesis in smaller epiblasts

To examine how apical actin polymerization could drive lumen opening, we first performed time-lapse imaging of fluorescently labelled actin during early lumenogenesis. Interestingly, lumen opening correlated with apical actin polymerization of only a few cells in the epiblast and, in some cases, specifically correlated with increase in apical length of a single cell while other cells maintained relatively constant apical lengths (Fig. 3, A and B, and Video S6). Though apical lengths are expected to increase with increasing lumen area, it was striking to see large cell-cell variations in growth dynamics for smaller epiblasts (Fig. 3, C and D). This heterogeneity implied that a cell-dependent mechanism drove lumenogenesis as a cell-independent mechanism, such as pressure, would be expected to increase the apical length of all cells. In line with these features, smaller epiblasts generated radially asymmetric matrix deformations (Fig. 3E).

**Figure 3.**
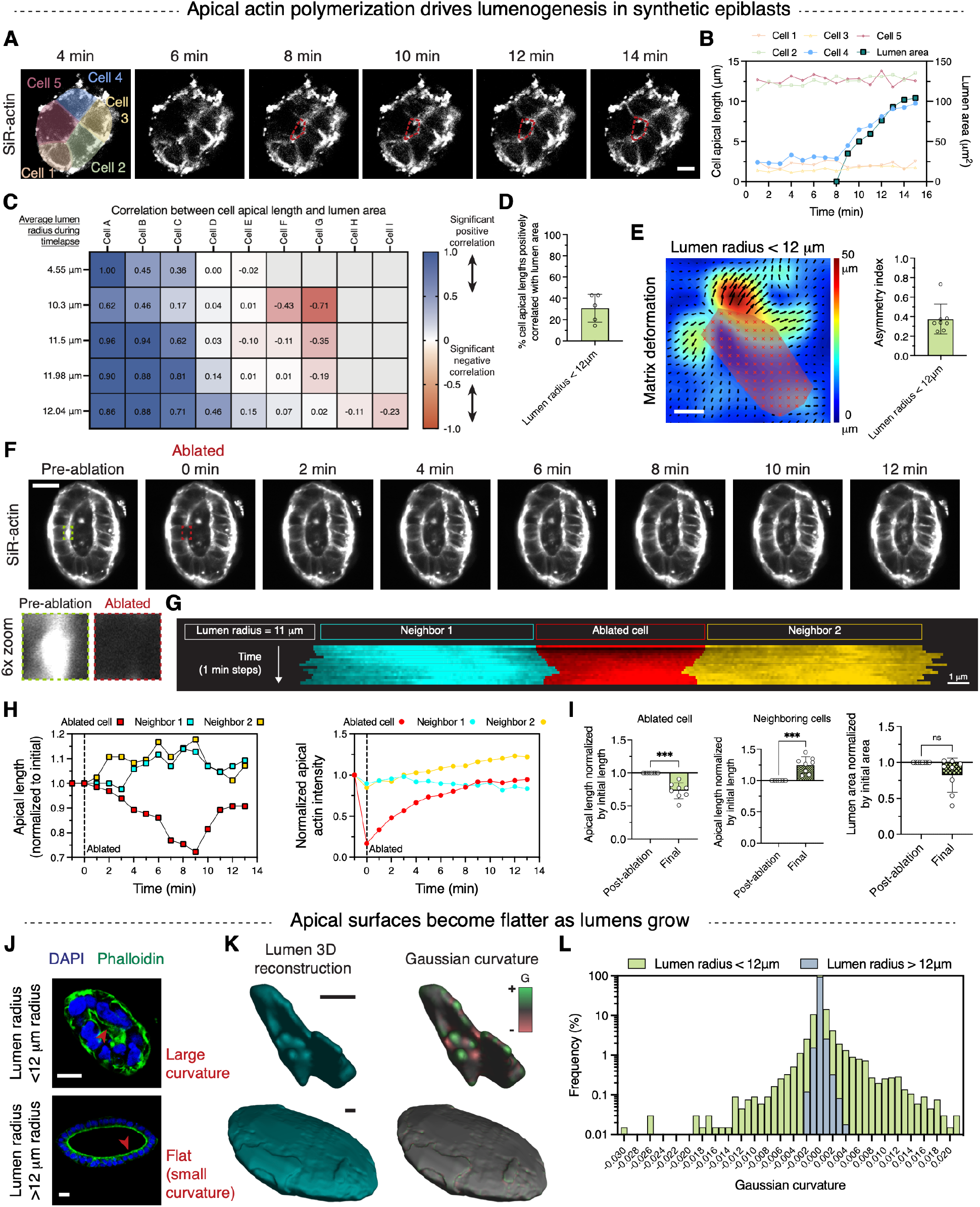
Apical actin polymerization drives lumenogenesis in smaller epiblasts. **(A)** Timelapse (min) images of actin during lumen opening. Red outlines indicate luminal surface. **(B)** Quantification of cell apical length and lumen area during lumen growth for epiblast shown in (A). **(C)** Spearman correlation values between lumen area and cell apical lengths for smaller epiblasts. Rows indicate each epiblast’s apical lengths correlated to corresponding lumen area. Spearman correlation values for each pair are listed and range between: 1 (perfect correlation), 0 (no correlation), and −1 (perfect anti-correlation). For Spearman correlation values > 0.5, p-value is < 0.05. Cells are listed in decreasing order of their respective Spearman correlation values. **(D)** Percent cells per epiblast whose apical lengths are positively correlated (Spearman correlation value > 0.5) with lumen area (mean ± s.d.). **(E)** Representative matrix deformations generated during lumen growth in smaller epiblasts and quantification of radial asymmetry in matrix deformation (mean ± s.d). Asymmetry index = magnitude of vector sum of deformations / average magnitude of deformations. **(F)** Apical actin ablation and recovery. Outlines indicate ablated surface. **(G)** Kymograph showing apical actin of ablated and neighboring cells for epiblast shown in (F). Length of colored lines = apical length; intensity of colored lines = average apical actin intensity. **(H)** Quantification of apical length and actin intensity of ablated and neighboring cells for epiblast shown in (F). **(I)** Quantification of post-ablation and final apical lengths of ablated and neighboring cells, and lumen area for smaller epiblasts. Apical lengths of the two neighbors were averaged (mean ± s.d.; ***p < 0.001, ns: not significant p > 0.05, Mann-Whitney; *n* = 8). **(J)** Representative cross-sectional immunostains of smaller and larger epiblasts. **(K)** 3D reconstruction of lumenal surface and quantification of Gaussian curvature for epiblasts shown in (J). **(L)** Frequency distribution of Gaussian curvature for epiblasts shown in (J). Scale bars: 10 μm (A and K), 20 μm (E, F and J), 1 μm (G).

Next, to test if apical actin polymerization in each cell can generate a pushing force on its neighbors, resulting in net hydrogel deformation, we performed laser ablation of apical actin in smaller epiblasts (Fig. 3F and Video S7). Remarkably, no immediate change in apical length was observed post ablation, suggesting that the apical actin mesh is neither under compression nor under tension (Fig. 3, F to I, Fig. S4, and Video S7). As the hydrogels are both viscoelastic and viscoplastic, this observation suggested that the stresses resisting the epiblast expansion were rapidly relaxed and the hydrogel was plastically deformed, so that residual stresses remaining on the epiblast are low at any given timepoint. Over a timescale of minutes following ablation, however, apical length of the ablated cell decreased, while that of neighboring cells increased, indicating active actin polymerization in the neighboring cells (Fig. 3, G to I, Fig. S4, and Video S7). But, as the apical actin in the ablated cell began to recover, the ablated cell pushed back on the neighboring cells, regrowing its apical length (Fig. 3, G and H, and Fig. S4). These ablation studies directly connect actin network growth to apical expansion and indicate the following interpretation. In smaller epiblasts, cells resist the growth of apical actin in neighboring cells and when such resistance is disrupted, say via ablation, actin in cells neighboring the ablated cell, can actively polymerize, increasing their apical lengths. Overall, these observations point to quasi-static actin growth where actin polymerization generated forces drive apical growth and lumen expansion, and stress relaxation and plasticity in the hydrogel prevent large tension or compression from building up in hydrogel and thus in the apical actin mesh as well.

Under the idea that actin polymerization at the apical surface of single cells drives lumen opening, cell-cell variations in actin polymerization rate and corresponding apical actin mechanics in smaller epiblasts, should result in a wide distribution of apical curvatures (Figs. 2J and 3, A to D). Analysis of lumen curvature showed a broad range of local curvatures in smaller lumens, as cells build their apical surfaces (Fig. 3, J to L). However, once all cells reach a mature apical size with a dense apical actin mesh, which is the case in larger epiblasts (Fig. 2, I and J), apical surfaces became uniformly flatter (Fig. 3, J to L). Taken together, these results show that apical actin polymerization at the apical surface of single cells drives lumen opening and expansion.

### Computational model of apical actin polymerization driven lumenogenesis

To better understand the physics of how apical actin polymerization can drive lumenogenesis, we developed a theoretical model and performed computational simulations. The model consisted of a cell cluster in an elastic hydrogel with a nascent lumen in the interior of the cluster. Cells actively polymerize apical actin and pump ions through the apical and basal surfaces. Water fluxes are driven by osmotic and hydrostatic pressure gradients that result from ion flux and pressure, but paracellular leaks can dissipate ion concentration gradients and allow water flux into or out of the lumen (Fig. 4A). In smaller clusters with leaky junctions, ion concentrations did not build up in the lumen and thus there was no difference in osmotic pressure between the lumen and the hydrogel (Fig. 4B). However, apical actin polymerization alone was sufficient to cause lumen opening and expansion, generating sufficient force to overcome the resistance of the hydrogel (Fig. 4, C to E). Longer apical lengths due to actin polymerization resulted in larger lumen sizes (Fig. 4C). Importantly, the model predicted that apical actin network stiffness was critical for driving lumen opening; reduced cortical stiffness led to smaller lumens (Fig. 4, D and E). Overall, our computational model demonstrates that osmotic pressure gradients cannot build up with sufficient paracellular leakage of ions, and that apical actin polymerization alone can drive lumen opening, implicating both actin polymerization forces and apical actin stiffness in mediating this connection.

**Figure 4.**
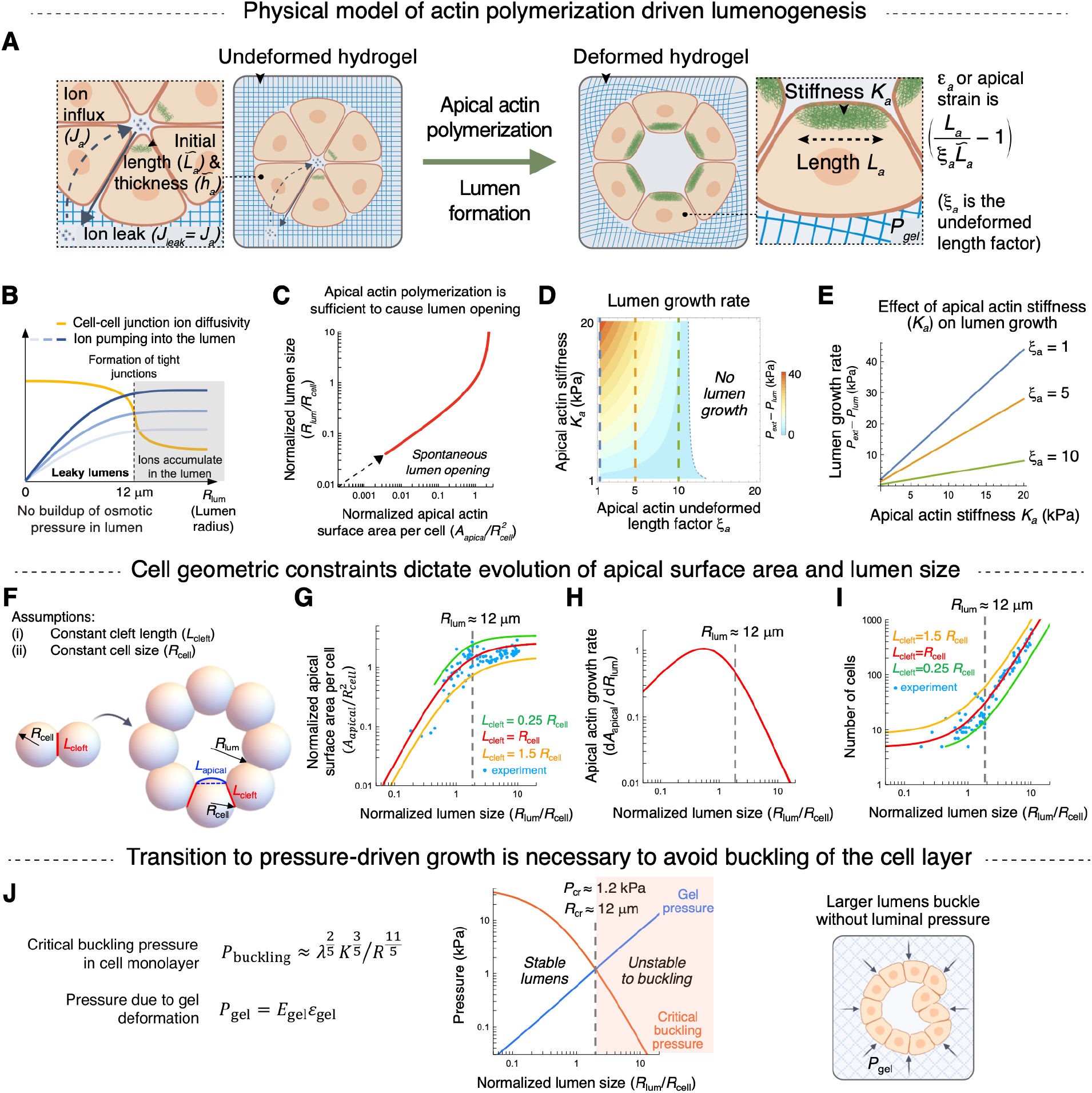
Physical model of actin polymerization driven lumenogenesis. **(A)** Schematic of the theoretical model of actin polymerization driven lumen growth. Hydrogel is considered linear elastic for this model with a modulus of 1 kPa which is the relaxed modulus of viscoelastic alginate hydrogels used in experiments. L̃*_a_*: initial length of apical actin mesh; *L_a_*: final length of apical actin mesh; *ξ*_a_: undeformed length factor; *K_a_*: stiffness of apical actin mesh; *J_a_*: ion flux into lumen; *J_leak_*: ion leak along the leaky junctions; *P_gel_*: stress exerted by the hydrogel on the cell cluster. **(B)** Model shows that osmotic pressure does not build in leaky lumens. Ions pumped into the lumen diffuse out along the leaky junctions in smaller epiblasts preventing buildup of osmotic pressure. **(C)** Cells actively polymerize apical actin that generates stress to deform the hydrogel and drive lumen opening. Dashed line indicates spontaneous lumen opening. Cell volume and cell thickness or cleft length are assumed to be constant. **(D and E)** Plots showing that increase in apical actin stiffness (*K_a_*) and decrease in undeformed length factor (*ξ*_a_) result in higher lumen growth rates. Lumen growth rates as a function of apical actin stiffness (*K_a_*) for given values of undeformed length factor (*ξ*_a_) are shown in (E). All other parameters were kept constant. *h_a_* = *h_b_* = 0.6 μm, *R_cell_* = 6 μm, *L_a_*/L̃*_a_* = 12, *L_b_*/L̃*_b_* = 1. **(F to I)** Schematic and predictions of a simple analytical model for estimating evolution of cell apical surface area and cell number with increase in lumen size, for fixed cell volume and cell thickness or cleft length. **(J)** Buckling pressure considerations predict a transition to pressure-driven lumen growth at ~12 μm lumen radius to prevent cell layer buckling.

We next sought to understand what factors necessitate rapid apical growth in smaller epiblasts and a transition to a different lumen growth mechanism in larger epiblasts. To answer this, we developed a simple analytical model to estimate lumen sizes and apical surface areas under two geometric constraints from experiments: (i) constant cell volume (or radius) (Fig. S3F), and (ii) constant cell thickness or cleft length (Fig. S4G and Fig. 4F). The resulting lumen sizes and apical lengths predicted by this model were in excellent agreement with the experimental observations, highlighting that rapid apical growth is necessary for lumen expansion in smaller epiblasts (Fig. 4, G to I). As lumens grow however, epiblasts become more susceptible to buckling (Trushko et al., 2020) and also stresses begin to accumulate in the hydrogel. Thus, there is a critical lumen size where pressure accumulated in the hydrogel becomes larger than the buckling pressure (Fig. 4J). For lumens to grow beyond this critical size, lumenal pressure is required to balance the hydrogel pressure and avoid buckling of the cell layer.

Overall, our computational model elucidates how apical actin polymerization drives lumen growth in smaller epiblasts, highlights geometric constraints that necessitate rapid apical growth in smaller epiblasts and predicts a transition to pressure driven growth in larger epiblasts to avoid buckling.

### Mechanism of lumen growth switches to osmotic pressure in larger epiblasts

As apical actin polymerization is stalled in larger epiblasts, we next examined the mechanism driving further growth of these lumens. Concomitant with the maturation of the apical surfaces (Fig. 2, I to L) at a lumen radius of ~12 μm, synthetic epiblasts form functional tight junctions (Fig. 1, G and H, and Fig. S2, A to E and H), which could allow osmotic pressure to build inside the lumen. Further, these lumens have convex, bulged out shapes (Fig. 1, L to N). Overall, these characteristics are consistent with osmotic pressure driven lumen growth (Chugh et al., 2022).

To test if osmotic pressure was responsible for growth of larger lumens, we performed time-lapse imaging. Unlike actin polymerization driven lumen growth, apical lengths of most cells in larger epiblasts generally increased over time and positively correlated with increase in lumen area (Fig. 5, A to D, and Video S8). Subsequently, larger epiblasts generated radially uniform matrix deformations as they grew (Fig. 5, E to G). Pressure-driven growth is governed by Young-Laplace law, which necessitates cells to be under tension to balance the luminal pressure (Chan et al., 2019; Latorre et al., 2018). Thus, laser ablation of apical actin was performed to examine if cells were under tension or compression. Ablated cells exhibited an immediate increase in apical length post-ablation revealing that cells were under tension (Fig. 5, H to K, and Video S9). With time, apical length of ablated cells increased further while no change in neighboring cells was observed (Fig. 5, H to K, and Video S9). Overall, these results provide strong evidence that osmotic pressure drives lumen growth in larger epiblasts.

**Figure 5.**
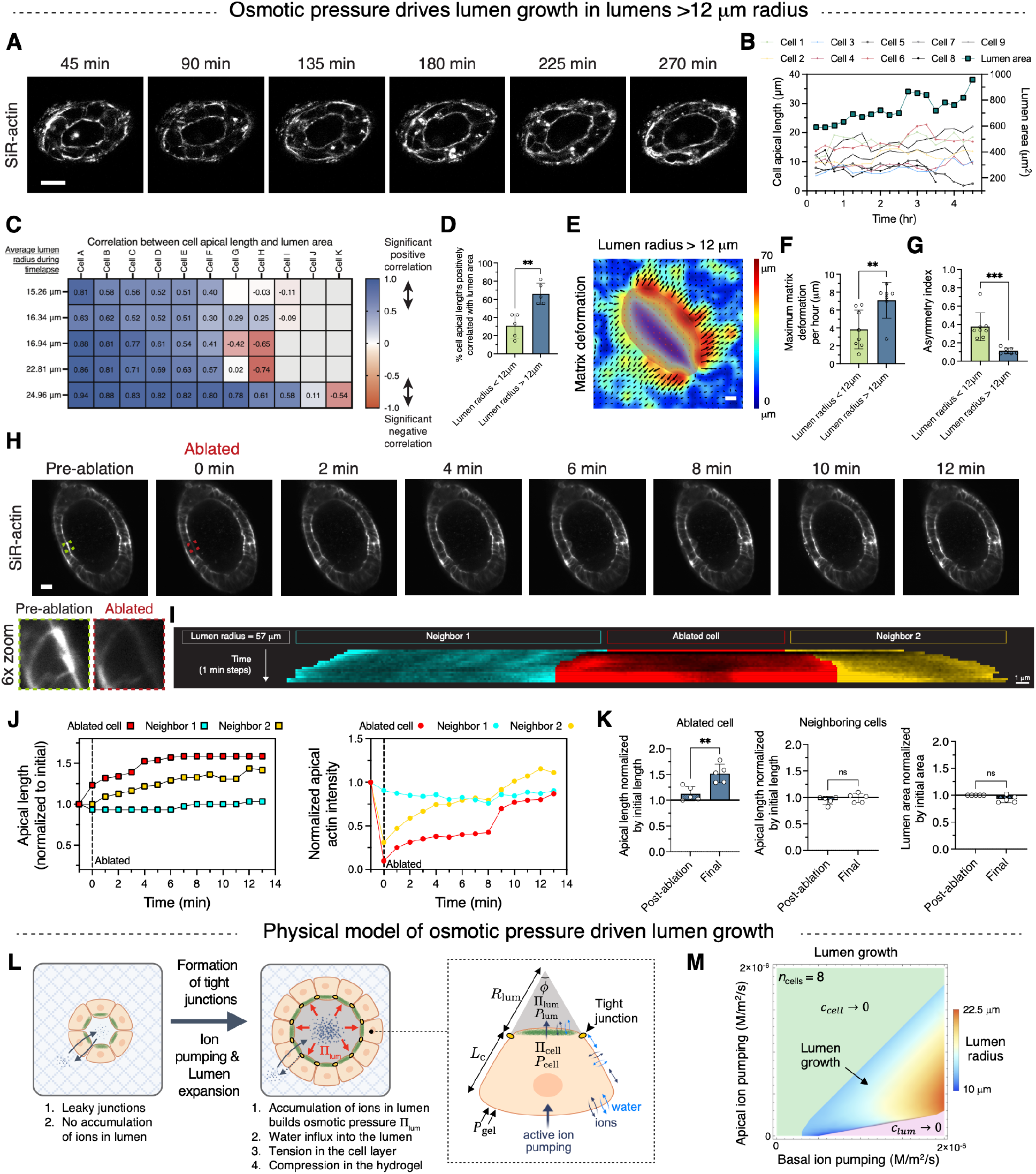
Osmotic pressure drives lumen growth in larger epiblasts. **(A)** Timelapse (min) images of actin during lumen opening. **(B)** Quantification of cell apical length and lumen area during lumen growth for epiblast shown in (A). **(C)** Spearman correlation values between lumen area and cell apical lengths for larger epiblasts. Rows indicate each epiblast’s apical lengths correlated to corresponding lumen area. Cells are listed in decreasing order of their respective Spearman correlation values. **(D)** Percent cells per epiblast whose apical lengths are positively correlated (Spearman correlation value > 0.5) with lumen area (mean ± s.d.; **p < 0.01, Mann-Whitney; *n* = 5 (smaller epiblasts), 5 (larger epiblasts)). **(E)** Representative matrix deformations generated during lumen growth in larger epiblasts. **(F)** Quantification of maximum matrix deformation per hour (mean ± s.d.; **p < 0.01, Mann-Whitney; *n* = 8 (smaller epiblasts), 7 (larger epiblasts)). **(G)** Quantification of radial asymmetry in matrix deformation (mean ± s.d.; ***p < 0.001, Mann-Whitney; *n* = 8 (smaller epiblasts), 7 (larger epiblasts)). Asymmetry index = magnitude of vector sum of deformations / average magnitude of deformations. **(H)** Apical actin ablation and recovery. Red outline indicates ablated surface. **(I)** Kymograph showing apical actin of ablated and neighboring cells for epiblast shown in (H). **(J)** Quantification of apical length and actin intensity of ablated and neighboring cells for epiblast shown in (H). **(K)** Quantification of post-ablation and final apical lengths of ablated and neighboring cells, and lumen area for larger epiblasts. Apical lengths of the two neighbors were averaged (mean ± s.d.; **p < 0.01, ns: not significant p > 0.05, Mann-Whitney; *n* = 5). **(L and M)** Schematic and predictions of the theoretical model of osmotic pressure driven lumen growth. Ion pumping into the lumen builds osmotic pressure as ions cannot diffuse out along tight junctions. *c_cell_* is the total concentration of ions in the cells at equilibrium and *c_lum_* is the total concentration of ions in the lumen at equilibrium. Scale bars: 20 μm (A, E and H), 1 μm (I).

To better understand the physics of osmotic pressure driven lumen growth, we applied our numerical model to larger epiblasts with tight junctions. In this case, ion pumping into the lumen increased osmotic pressure and resulted in robust lumen growth (Fig. 5, L and M). Predicted lumen sizes were in excellent agreement with experimental observations (Fig. S5). Balance between apical and basal ion pumping was found to be critical for buildup of osmotic pressure (Fig. 5M). Higher basal pumping without apical pumping, and vice versa, prevented accumulation of ions in the lumen and no lumen growth occurred (Fig. 5M). Overall, our experiments and model indicate that ion pumping in the presence of tight junctions builds osmotic pressure to drive the growth of larger lumens.

### Human epiblasts upregulate actin polymerization related genes during lumenogenesis

Finally, to investigate the in vivo relevance of the mechanisms discovered in synthetic epiblasts, we analyzed transcriptional signatures of the human epiblast as a function of developmental time using single cell RNA sequencing data generated from peri-implantation human embryos (Xiang et al., 2020; Zhou et al., 2019). Epiblast lumen forms soon after implantation at ~7 d.p.f and expands in volume up to gastrulation at ~14 d.p.f (Fig. 1, A and B) (Shahbazi and Zernicka-Goetz, 2018). Analysis of epiblast cells from 7 to 14 d.p.f using KNN (k-nearest neighbor) based clustering of UMAP (Uniform Manifold Approximation and Projection) dimensionality reduction plots revealed two cell subpopulations (Fig. 6, A and B). As the two epiblast subpopulations were roughly separated based on developmental time, we annotated these as early and late epiblasts (Fig. 6B). As expected, early epiblast cells showed higher expression of naïve pluripotency markers such as *DNMT3L* and *KLF4* and lower expression of primed pluripotency marker *SFRP2*, as compared to late epiblast cells (Fig. 6, C and D). Interestingly, several actin polymerization related genes including those encoding for proteins in the Arp2/3 complex such as *ARPC1B, ARPC5, ARPC2* were upregulated in the early epiblast but transitioned to a lower expression level in the late epiblast (Fig. 6E). This is consistent with the expectation from our synthetic epiblast findings where cells actively build a branched apical actin network at earlier stages of lumenogenesis but transition to an equilibrium apical size at later stages (Fig. 2I). Similar transcriptional signatures were observed in a different single cell RNA sequencing dataset (Zhou et al., 2019) of 8-12 d.p.f human embryos as well (Fig. S6). Altogether, the upregulation of actin polymerization genes early in lumenogenesis are suggestive that actin polymerization may drive early lumen formation in the human epiblast, as we have found in synthetic epiblasts in this study.

**Figure 6.**
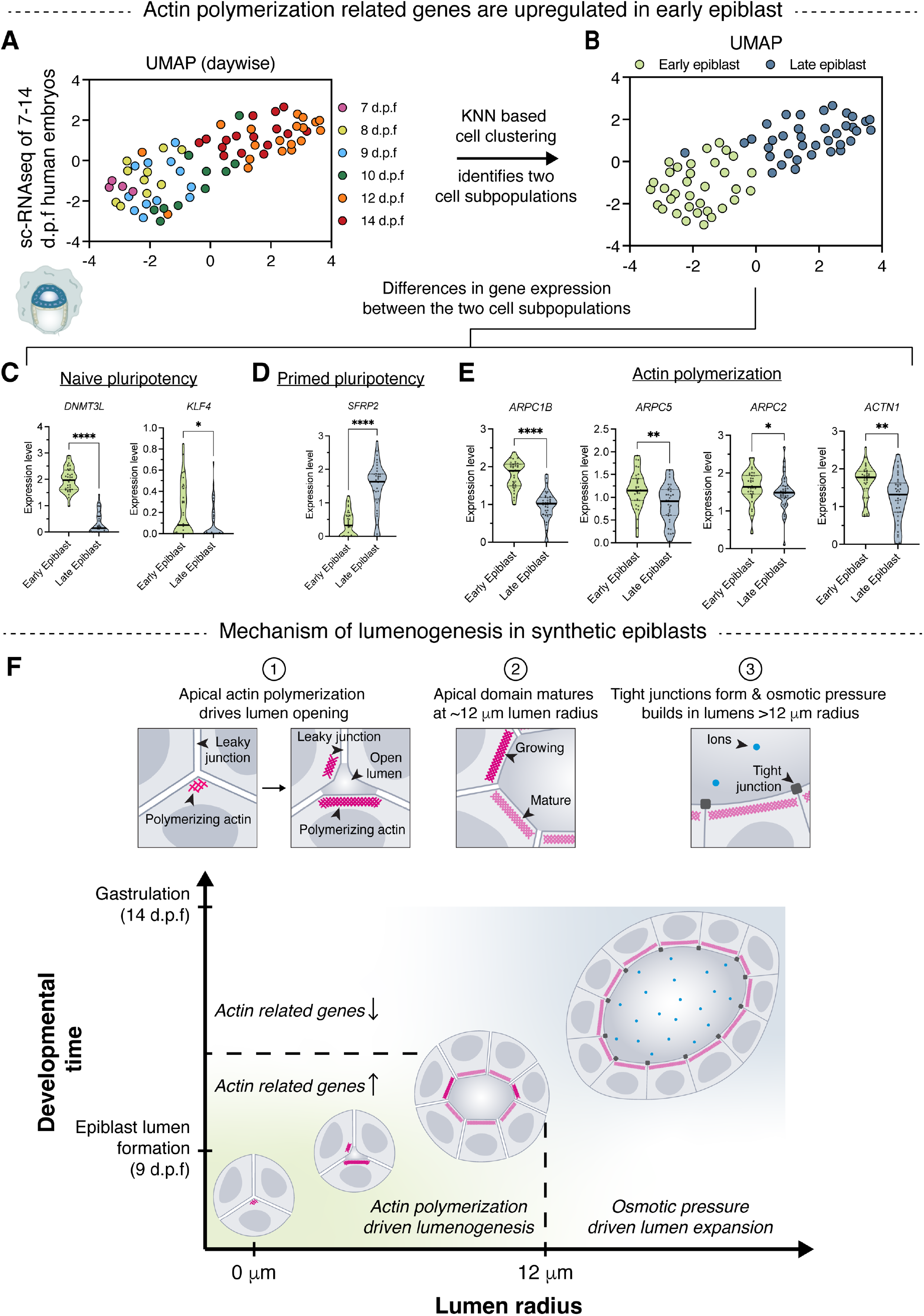
Human epiblasts show transcriptional signatures consistent with actin polymerization driven lumenogenesis. **(A)** UMAP plot of human epiblast cells generated from scRNA-seq data of peri-implantation human embryos (Xiang et al., 2020) (71 epiblast cells) colored with timepoints (d.p.f). **(B)** KNN (k-nearest neighbor) based cell clustering. The two subpopulations are annotated as early and late epiblast. **(C and E)** Normalized expression level of naïve pluripotency genes (C), primed pluripotency genes (D) and actin polymerization related genes (E) (median and quartiles; ****p < 0.0001, **p < 0.01, *p < 0.05, Mann-Whitney; *n* = 34 (early epiblast cells), 37 (late epiblast cells)). **(F)** Summary of mechanisms driving lumenogenesis in human epiblasts.

## Discussion

Taken together, our experimental and simulation results uncover the physical mechanisms of lumenogenesis in synthetic epiblasts. Synthetic epiblasts closely model the morphology and phenotype of human epiblasts. We describe two distinct lumen-size-dependent mechanisms that generate the force necessary for lumen expansion. Synthetic epiblasts with lumens smaller than ~12 μm radius lack functional tight junctions, allow free diffusion of ions and macromolecules between the lumen and the hydrogel, and prevent large osmotic pressures from building up. Lumen growth in these smaller epiblasts is driven by actin polymerization into a dense network on the apical surface via N-WASP, Arp2/3 and formins. Force generation by actin polymerization, aided by rapid stress relaxation in the hydrogel, ultimately drives lumen growth and overall expansion of the synthetic epiblast in the hydrogel (Fig. 6F). When apical actin mesh in individual cells reaches a defined equilibrium size at a lumen radius of ~12 μm, coinciding with the formation of tight junctions, the mechanism of lumen growth switches to osmotic pressure gradient driven (Fig. 6F). Lastly, transcriptional expression profiles of human epiblasts support the existence of similar mechanisms in vivo as those discovered in synthetic epiblasts.

Mechanisms of human epiblast lumenogenesis have been difficult to explore owing to ethical concerns, technical challenges with in vitro culture of human embryos and limitations of stem-cell models of the human epiblast (Carleton et al., 2022). Matrigel or reconstituted basement membrane dependent models of the human epiblast provide a limited window into lumen growth as cells quickly differentiate in culture (Indana et al., 2021; Simunovic et al., 2019), thus only allowing the study of early polarization and lumen growth events (Taniguchi et al., 2017; Taniguchi et al., 2015; Wang et al., 2021). In mice, while peri-implantation development is significantly different from humans, a few different mechanisms have been suggested to play a role in epiblast lumenogenesis: (i) electrostatic repulsion between apically deposited anti-adhesive proteins such as podocalyxin which have a high negative charge (Bedzhov and Zernicka-Goetz, 2014) and (ii) luminal fluid transport due to osmotic gradients (Kim et al., 2021). However, these mechanisms alone fail to provide sufficient physical explanations for sustained lumen expansion. While anti-adhesive proteins such as podocalyxin can help create a “non-stick” apical surface, electrostatic repulsive forces are negligible on a length scale of microns due to Debye-Hückel screening in electrolyte solutions (Torres-Sánchez et al., 2021). Directed fluid transport on the other hand does not in itself generate force for lumen expansion unless the fluid is pressurized or accompanied by other cellular force generating mechanisms. By using synthetic hydrogels which provide a prolonged window into human epiblast lumen formation and expansion, we unraveled the physical force generating mechanisms responsible for sustained lumen growth.

Actin polymerization and assembly into branched networks via Arp2/3 complex is known to generate pushing forces and drive several cellular processes including lamellipodial protrusions during migration, vesicle trafficking and polarization (Gautreau et al., 2022). Such actin structures also play a vital role during early embryogenesis (Lim and Plachta, 2021). For example, expanding apical actin rings in a pre-implantation mouse embryo push cells against each other, stabilizing cell-cell junctions (Zenker et al., 2018) and allowing the formation of a pressurized blastocyst cavity (Chan et al., 2019). Here, we discovered a novel mechanism of actin-polymerization-driven lumen formation in the human epiblast. Human epiblast tissue is confined by extraembryonic cells and the pressurized blastocyst cavity (~1 to 2 kPa pressure (Chan et al., 2019)) and thus epiblast lumenogenesis requires force generation. While actin structures are susceptible to buckling, we find that apical actin polymerization and formation of a branched actin mesh generates sufficient force to drive epiblast lumen opening. Apical actin as well as actomyosin networks are a common feature of epithelia, but the structural features of apical actin mesh and how actin polymerization is physically aligned to drive lumenogenesis are yet to be explored.

The existence of two distinct mechanisms of lumen growth, dependent on a critical lumen size at which tight junction formation and apical maturation occur, highlights close crosstalk between the cell polarity machinery, growth of apical domains, tight junctions, and lumen size. Cell geometries are also tightly controlled with cell volume and thickness staying roughly constant during both actin polymerization and osmotic pressure driven lumen growth. Further, cell-layer stretching and cycles of lumen inflation and collapse, which are characteristic of pressure driven lumen growth in other model systems (Chan et al., 2019; Latorre et al., 2018; Mosaliganti et al., 2019; Ruiz-Herrero et al., 2017), are not observed in synthetic epiblasts, suggesting that pressures generated in the epiblast are relatively low. While high pressures are useful for disrupting structures such as zona pellucida during blastocyst hatching (Kim and Bedzhov, 2022), they could cause tissue rupture (Chan et al., 2019; Latorre et al., 2018) and compromise embryo integrity. Moreover, lumen volumes and cell numbers closely follow a power law relationship during pressure driven growth, highlighting a careful balance between pressure magnitude and cell number, thereby allowing cells to maintain a fixed volume and thickness. Such control over embryo size could be pivotal for subsequent embryonic patterning events such as amnion formation and gastrulation by establishing appropriate signaling gradients (Kim et al., 2021; Ryan et al., 2019; Zhang et al., 2019). Notably, we did also find mechanisms of robustness in this system. While inhibition of actin polymerization disrupts early lumen formation, lumens ultimately form as cells multiply even with continuous actin inhibition, highlighting the presence of alternate mechanisms of epiblast lumenogenesis to ensure robust development, though the delay in lumenogenesis could possibly impact other developmental processes. Overall, our results provide a quantitative understanding of the mechanisms that drive epiblast lumenogenesis and suggest that size control and robustness are inherently coded into these mechanisms.

Though synthetic epiblasts replicate many features of the human epiblast, a few key differences remain. For example, our epiblast model does not undergo amnion formation, an event that coincides with epiblast cavity growth. Moreover, our model lacks extraembryonic cells and their corresponding biochemical signals. The role of extraembryonic cells and mechanisms of amnion formation require further investigations. Also, several new questions arise about the function of the epiblast lumen. While the epiblast lumen has been suggested to shield the epiblast cells from extraembryonic signaling to ensure robust gastrulation (Pfeffer, 2022), the existence of size-dependent mechanisms of lumen growth points to a larger role for the epiblast lumen in regulating embryo size and orchestrating embryonic development. To our knowledge, the actin polymerization driven lumenogenesis mechanism discovered here is the only force-generating, pressure-independent lumenogenesis mechanism and could be at play in other model systems as well. In conclusion, this study advances our understanding of human embryonic development and expands our knowledge of the biological toolkits that cells utilize to make a lumen.

## Acknowledgments

The authors gratefully acknowledge all current and past members of the Chaudhuri Lab for helpful discussions. The authors acknowledge the Stanford Cell Sciences Imaging Facility (CSIF) for microscope (Zeiss lattice light sheet 7 and Zeiss LSM 780 inverted multiphoton laser scanning confocal microscope) and software access (Imaris). The authors acknowledge Gordon Wang, Ph.D., and the Stanford Neuroscience Microscopy Service for access to and help with super-resolution microscopy (Zeiss Airyscan2 LSM 980 inverted confocal microscope). The authors thank Zeiss specialists Win Zaw, Ph.D. and Benjamin Figueroa, Ph.D. for help with joint deconvolution (jDCV) processing of Airyscan2 super-resolution images. The authors thank Dr. Vittorio Sebastiano (Department of Obstetrics and Gynecology, Stanford University) for providing the hiPSC line derived from BJ fibroblasts and Dr. Marc Levenston (Department of Mechanical Engineering, Stanford University) for use of mechanical testing equipment (Instron). This work was supported by a Stanford School of Engineering Graduate Fellowship to D.I., a Stanford Bio-X Interdisciplinary Initiatives Program Seed grant for O.C. and N.B., and a National Science Foundation grant MCB 2148041 for O.C.

## Author contributions

D.I. and O.C. conceived and designed the experiments. D.I. performed the experiments, data analysis, and statistical tests. D.I. and Y.L. performed experiments using light sheet microscopy. A.Z. and V.B.S performed computational simulations and analysis. N.B. and A.R.D. contributed to experimental design and provided analytical guidance. D.I., A.Z., V.B.S. and O.C. wrote the manuscript with input from all authors.

## Data and materials availability

All data generated or analyzed during this study are included in the main manuscript and supplementary materials. Raw data and materials for all data are available from the corresponding author upon reasonable request.

## Code availability

Custom scripts of code for the computational models, analysis of single cell RNA-sequencing datasets and image analysis are available from the corresponding author upon reasonable request.

## Ethics statement

The synthetic epiblasts used in this study lack extraembryonic cells including hypoblast and trophectoderm, and do not have human organismal potential, i.e., cannot form a whole human embryo. Furthermore, all experiments were terminated by no later than day 7 of culture. All protocols used in this work with hiPSCs have been approved by the Stanford University Stem Cell Research Oversight Committee (SCRO protocol 837).

## Materials and methods

### Alginate preparation

Sodium alginate rich in guluronic acid blocks was purchased (ProNova UP VLVG; 28 kDa molecular weight; Dupont). RGD (arginine–glycine–aspartate) peptides were coupled to alginate using carbodiimide chemistry (Rowley and Mooney, 2002). First, alginate was dissolved overnight at 1% (w/v) in a 0.1 M MES hydrate (Sigma-Aldrich M8250), 0.3 M sodium chloride (Fisher Scientific S671) buffer with a pH of 6.5. *N*-hydroxysulfosuccinimide (Sulfo-NHS, Thermo Fisher Scientific 24510), *N*-(3-dimethylaminopropyl)-*N*’-ethylcarbodiimide hydrochloride (EDC, Sigma-Aldrich E6383) and GGGGRGDSP (Peptide 2.0) peptide were sequentially mixed in the alginate solution and the reaction was allowed to proceed for 20 hr until quenched by adding hydroxylamine hydrochloride (Sigma-Aldrich 255580). The alginate was then dialyzed in deionized water for 3 days, purified with activated charcoal, sterile filtered, frozen, lyophilized and stored at −20°C. For cell encapsulation, lyophilized alginate was reconstituted at 3% (w/v) in serum-free DMEM/F-12 (Thermo Fisher Scientific 11330057). Reconstituted 3% (w/v) alginate was diluted with solution containing cells and crosslinked using calcium sulfate to make hydrogels with 2% (w/v) final alginate concentration, initial elastic modulus of 20 kPa, loss tangent of ~0.08, stress relaxation half-time of ~70 s, and 1500 μM RGD density as described previously (Indana et al., 2021). Detailed protocols for RGD-conjugation of alginate, preparation of alginate hydrogels and exact recipes for alginate hydrogels used in this study have been published previously (Charbonier et al., 2021; Indana et al., 2021).

### Hydrogel mechanical characterization

Compression tests were performed using an Instron 5848 MicroTester to quantify the initial elastic modulus and stress relaxation behavior of the alginate hydrogels. Alginate disks of 2 mm thickness and 4 mm diameter were prepared and equilibrated in DMEM/F-12 for 24 hr. Unconfined compression tests were then performed on alginate disks using a 4 mm diameter cylindrical probe. Gels were compressed from 0 to 10% compressive strain at a deformation rate of 1 mm per min. 10% compressive strain was then maintained for 1 hr and the corresponding stress was measured over time (stress relaxation test). To calculate the initial elastic modulus, a straight line was fitted to stress vs. strain data for the initial strain ramp between 5% and 10% compressive strain. The slope of this linear fit was reported as the initial elastic modulus. Next, to quantify the stress relaxation behavior at 10% compressive strain, the time at which relaxation modulus drops to half of its initial value was measured and reported as τ_1/2_. Final relaxed modulus during the stress relaxation test (~1 kPa) was taken as the effective hydrogel stiffness for cellular processes (including lumen growth) that were much slower than the hydrogel stress relaxation half-time (~70 s).

### Cell lines and culture

Two hiPSC lines were used in this study. First, was a hiPSC line (RiPSC.BJ) generated through synthetic mRNA reprogramming of BJ human fibroblast cells (Durruthy-Durruthy et al., 2014) (a gift from Dr. Vittorio Sebastiano (Department of Obstetrics and Gynecology, Stanford University)). Second, was a hiPSC line purchased from Coriell Institute (AICS-0024) in which MYH10 has been endogenously tagged with mEGFP using CRISPR/Cas9 technology in WTC-11 (GM25256) hiPSCs. hiPSCs were first expanded to generate a large cell bank within 2 passages of purchased or gifted cells: passage 29 and 30 for MYH10-mEGFP hiPSCs and passage 16 and 17 for untagged hiPSCs. Both hiPSC lines have been authenticated by original sources for successful differentiation to the three germ layers and also authenticated in-house for pluripotency by Oct4, Sox2 and Nanog immunostaining.

hiPSCs were cultured on TC-treated 100 mm dish (Corning 430167) coated with LDEV-free hESC-qualified Matrigel (Corning 354277) in mTeSR1 media (STEMCELL Technologies) at 37°C in 5% CO_2_. hiPSCs cultured in mTeSR1 were used for encapsulation in alginate hydrogels at 70% confluency as single cells using Accutase (STEMCELL Technologies) following the manufacturer’s protocol. For each experiment, a new hiPSC vial (of passage numbers listed above) was thawed, cultured as described above and encapsulated in alginate hydrogels without continued passaging. This was done to ensure high-quality of hiPSCs and to maintain hiPSCs at a low passage number and normal karyotype as characterized previously (Durruthy-Durruthy et al., 2014). Both hiPSC lines were checked for mycoplasma contamination and tested negative (LookOut Mycoplasma PCR Detection Kit, Sigma-Aldrich MP0035).

### Encapsulation of cells within hydrogels

To make hydrogels, appropriate volumes of 3% (w/v) alginate and dissociated single hiPSCs were added to a luer lock syringe (Cole-Parmer). In a second syringe, appropriate volumes of calcium sulfate and serum-free DMEM/F-12 were added. The two syringes were connected with a coupler and the solutions were mixed by passing them back and forth six times. The mixture of cell, alginate and calcium sulfate solution were either directly deposited into an 8-well Lab-Tek chamber slide (Thermo Fisher Scientific) or onto a hydrophobic glass plate which was then covered with another glass plate with a 1 mm spacer between plates. The cell alginate mixture was then allowed to gel for 30 mins at room temperature. Hydrogels had a final alginate density of 2% (w/v), cell concentration of 1 million per mL of hydrogel and calcium concentration of 33 mM. Hydrogels were punched out using a 6 mm diameter biopsy punch, immersed in mTeSR1 media with 10 μM ROCK inhibitor (Y-27632, STEMCELL Technologies) to prevent dissociation- induced apoptosis and transferred to an incubator at 37°C and 5% CO_2_. 24 hr post encapsulation, media was changed to mTeSR1, which was replenished daily.

### Sample preparation for immunofluorescence

hiPSCs cultured in hydrogels were fixed with 4% paraformaldehyde (PFA, Alfa Aesar) for 1 hr, and then washed three times with PBS containing Ca^2+^ and Mg^2+^ (cPBS, GE Healthcare). For immunostaining of whole synthetic epiblasts, these hydrogels were immersed in cPBS and stored at 4°C until used. For immunostaining two-dimensional sections of hydrogels, gels were placed in 30% (w/v) sucrose (Fisher Scientific) overnight and then transferred to 50% (v/v) of 30% (w/v) sucrose and OCT compound solution (Tissue-Tek) for 5 hr. The hydrogel was then embedded in OCT, frozen, and sectioned at 40 μm thickness using a cryostat (Leica CM1950).

### Immunofluorescence of two-dimensional sections and whole synthetic epiblasts

For two-dimensional sections, samples were washed three times in DPBS (Thermo Fisher Scientific), permeabilized with 0.5% Triton X-100 (Sigma) in DPBS for 15 mins and blocked for 1 hr with 1% bovine serum albumin (BSA, Sigma), 10% goat serum (Invitrogen), 0.3 M glycine (Fisher), and 0.1% Triton X-100 in DPBS. The samples were incubated overnight with primary antibodies. After washing three times with 0.1% Triton X-100 in DPBS, DAPI (1:1000 dilution; 5 mg/ml stock solution, Sigma-Aldrich) and Alexa Fluor 488 Phalloidin (1:80 dilution; Invitrogen A12379) were incubated for 1.5 hr to stain the nucleus and F-actin along with appropriate secondary antibodies, such as goat anti-Rabbit IgG AF647 (Invitrogen, 1:200 dilution) and goat anti-mouse IgG1 AF555 (Invitrogen, 1:200 dilution). Following three washes with 0.1% Triton X-100 in DPBS, ProLong Gold antifade reagent (Life Technologies) was applied to minimize photobleaching. Images were acquired on a Leica SP8 laser scanning confocal microscope using a HC PL APO 63×/1.4 NA oil immersion objective. Primary antibodies used in study were Oct4 (Cell Signaling Technology #2750), Sox2 (Cell Signaling Technology #3579), Nanog (Cell Signaling Technology #4903), Otx2 (R&D systems AF1979), Ezrin (Sigma-Aldrich E8897), Podocalyxin (R&D systems MAB1658), ZO-1 (Thermo Fisher Scientific 33-9100), phospho- myosin light chain 2 (Cell Signaling Technology #3674), Arp2/3 complex (Sigma-Aldrich MABT95), N-WASP (Thermo Fisher Scientific PA5-52198); all used at 1:200 dilution.

For immunostaining of whole synthetic epiblasts, PFA-fixed synthetic epiblasts were first recovered from alginate hydrogels by incubating the hydrogel for 5 mins at room temperature in 50 mM EDTA (ethylenediaminetetraacetic acid) solution. EDTA chelates calcium ions and dissolves the alginate hydrogel, following which mild centrifugation was performed to collect the synthetic epiblasts. Next, for immunostaining, the same protocol as above was followed with slight modifications: (i) all steps were performed in ultra-low attachment 24-well plates (Corning), (ii) longer incubation – 1 hr permeabilization, 3 hr blocking and overnight secondary antibody steps. Images were acquired on a Leica SP8 laser scanning confocal microscope using a HC FLUOTAR L 25×/0.95 NA water immersion objective.

### SYTOX staining

hiPSCs in alginate hydrogels were incubated with SYTOX Green Nucleic Acid Stain (1 μM; Invitrogen, Thermo Fisher Scientific S7020) for 1 hr at 37°C and imaged on a Leica SP8 laser scanning confocal microscope with a HC FLUOTAR L 25×/0.95 NA water immersion objective.

### Tight junction permeability studies

Cell impermeable fluorescent dextran was used to assess tight junction permeability in synthetic epiblasts. Dextrans with the following molecular weights were used: 3 kDa (Thermo Fisher Scientific D3328), 10 kDa (Thermo Fisher Scientific D1828), 40 kDa (Thermo Fisher Scientific D1829) and 70 kDa (Thermo Fisher Scientific D1830). All these dextrans are conjugated with Texas Red fluorophore and are zwitterionic. For permeability assay, dextran dissolved in mTeSR1 at a final concentration of 10 μM was added to the hydrogel and incubated at 37°C for 1 hr. Synthetic epiblasts were then imaged using Leica SP8 laser scanning confocal microscope with a HC FLUOTAR L 25×/0.95 NA water immersion objective.

To quantify tight junction permeability, outlines were drawn manually around the lumen boundary, inside a cell and in the hydrogel using both the fluorescent dextran and brightfield images to measure dextran intensity in the lumen, cell, and hydrogel using ImageJ (NIH). Cell dextran intensity normalized to that in the hydrogel was consistently ~0.1 confirming that dextrans were cell impermeable. Lumenal dextran intensity was then normalized to that in the hydrogel, called normalized lumenal dextran intensity (NLDI), as a measure of tight junction permeability. Lumen size and shape metrics were measured in ImageJ. Lumen radius was determined from measured lumen area assuming a perfect circle.

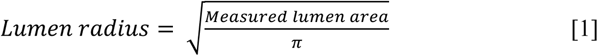

### Fluorescence recovery after photobleaching (FRAP) studies

For quantifying diffusion dynamics in smaller epiblasts which lacked tight junctions, fluorescence recovery of dextran was observed after photobleaching on a Zeiss LSM 780 laser scanning confocal microscope using a LCI PLAN NEO 25×/0.8 NA oil immersion objective at 37°C and 5% CO_2_. hiPSCs were incubated with 3 kDa dextran for 1 hr before imaging as described in the previous section. For photobleaching, lumen boundary was manually outlined and excited with a micro-point laser at 594 nm (as excitation peak of Texas Red-labelled-dextran is 595 nm) to bleach lumenal dextran using 100 scan iterations of 100% laser power (max power: 0.194 mW). Images were taken before and after bleaching, at 1 min intervals.

Recovery profiles obtained after photobleaching were measured as NLDI (see previous section) and normalized such that initial NLDI was 1 and post-bleach NLDI was 0. The data was then fit to a previously described experimental recovery curve which assumes bleaching of a 2D circular spot followed by free diffusion of non-bleached molecules into the bleached spot from all directions (Soumpasis, 1983):

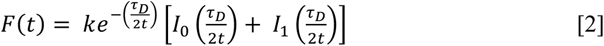

where *I*_0_ and *I*_4_ are modified Bessel functions of the first kind of zero and first order, k is the mobile fraction and *τ*_5_ is the characteristic diffusion time. τ_1/2_ (diffusion half-time) was calculated as the time at which fluorescence recovers to half the final equilibrated value. While this model accurately fit the experimental data (R^2^ > 0.99), the assumptions of this model are not completely valid for lumenal bleaching as dextran can diffuse into the lumen only via intercellular spaces and not along all radial directions. Thus, the diffusion coefficient obtained from *τ*_5_ cannot be directly compared to diffusion coefficients predicted by the Stokes-Einstein equation.

### Inhibition studies

For all small-molecule inhibition studies, the drug was added 24 hr before imaging or continuously starting on day 2 of culture (for plots labelled continuous inhibition). The inhibitors used were Blebbistatin (10 µM; Abcam ab120425, myosin II inhibitor), ML-7 (10 µM; Tocris Bioscience 4310, myosin light chain kinase inhibitor), ML-141 (2 µM; Tocris Bioscience 4266, selective Cdc42 Rho family inhibitor), Y-27632 (10 µM; STEMCELL 72304, ROCK inhibitor), CK-666 (50-100 µM; Sigma-Aldrich SML0006, Arp2/3 inhibitor), SMIFH2 (10-20 µM; Sigma-Aldrich S4826, formin inhibitor) and Wiskostatin (5-10 µM; Abcam ab141085, N-WASP inhibitor). All drugs were dissolved in dimethylsulfoxide (DMSO) and diluted in mTeSR1 media before adding to hiPSCs. DMSO alone was added to mTeSR1 media as a vehicle control. Percent clusters with lumen was manually quantified from brightfield images using ImageJ.

### SiR-actin staining and live cell imaging

For live imaging of actin in synthetic epiblasts, SiR-actin (100 nM; Cytoskeleton Inc. CY-SC001) was added to hiPSCs in hydrogels for 12 hr following manufacturer’s protocol. For timelapse imaging of actin, mTeSR1 media was supplemented with 100 nM SiR-actin (to maintain strong actin fluorescence) and 2.5 mM *N*-acetyl-L-cysteine (NAC, Sigma-Aldrich A9165; antioxidant to reduce phototoxicity). Fluorescent actin (excitation peak of SiR-actin is 652 nm) and brightfield images were acquired every 15 min for a total duration of 8 hr on a Leica SP8 laser scanning confocal microscope with a HC FLUOTAR L 25×/0.95 NA water immersion objective at 37°C and 5% CO_2_.

For quantifying timelapse data, lumen areas and apical lengths of individual cells were manually measured using ImageJ. To understand the correlation between lumen growth and increase in cell apical actin lengths for a fixed number of cells, 4 hr time windows were picked from acquired data, during which there was no change in cell number in both smaller and larger epiblasts. Correlation between lumen area and individual cell apical lengths was quantified by calculating Spearman correlation values between each pair of curves in GraphPad Prism (9.3.1). Percent cell apical lengths positively correlated with lumen area was calculated by counting number of cells whose correlation with lumen area has a Spearman value >0.5 and dividing by total number of cells in a given epiblast. 3D timelapse imaging of synthetic epiblasts with high z-resolution was not possible due to phototoxicity effects upon increased laser exposure.

### Quantification of hydrogel deformations

For quantifying hydrogel deformations, 1 µm diameter fluorescent carboxylate-modified microspheres (FluoSpheres, Thermo Fisher Scientific F8816) were encapsulated in alginate hydrogels. Timelapse images of fluorescent beads were collected during lumen growth in both smaller and larger epiblasts on a Leica SP8 laser scanning confocal microscope with a HC FLUOTAR L 25×/0.95 NA water immersion objective at 37°C and 5% CO_2_. Acquired images were corrected for drift using an ImageJ plugin (StackReg). Next, the drift-corrected images were used to calculate matrix deformations in MATLAB by tracking beads using a particle image velocimetry algorithm (PIVlab; open source code) using three cross-correlation windows (128 × 128, 64 × 64, and 32 × 32 pixel interrogation windows). Maximum matrix deformation was selected from within ~100 µm^2^ around the cells. Radial asymmetry of matrix deformations (asymmetry index) was quantified by taking the ratio of magnitude of vector sum of deformations to average magnitude of deformations. Finally, note that the direct estimation of forces from matrix deformations is challenging due to hydrogel viscoelasticity and plasticity. Thus, in this study, matrix deformations were used as a proxy for force generation, as matrix deformation only occurred as a result of cellular forces in these gels.

### Super-resolution microscopy

Super-resolution microscopy was performed to visualize apical actin mesh in SiR-actin stained synthetic epiblasts, using a Zeiss Airyscan2 LSM 980 inverted confocal microscope with a LCI PLAN NEO 25×/0.8 NA oil immersion objective at 37°C and 5% CO_2_. Images were acquired in super-resolution mode with a voxel size of 0.0974 × 0.0974 × 0.81 µm^3^ (x y z) and processed using 15 iterations of a 3D iterative joint deconvolution (jDCV) algorithm (Zeiss).

### Laser ablation studies

Laser ablation was performed to determine the stress state of actin in SiR-actin stained synthetic epiblasts, using a Zeiss LSM 780 laser scanning confocal microscope with a LCI PLAN NEO 25×/0.8 NA oil immersion objective at 37°C and 5% CO_2_. A micro-point laser at 405 nm was used to ablate apical actin or to cut through the entire thickness of a cell in both smaller and larger epiblasts using 150 scan iterations of 100% laser power (max power: 1.85 mW). Images were taken before and after ablation, at 1 min intervals.

To measure recoil or retraction of apical actin in synthetic epiblasts, apical lengths and apical actin intensity of ablated cell and its neighboring cells as well as lumen area were manually measured using ImageJ at each time-point: pre-ablation, immediately post-ablation and at 1 min intervals post-ablation till 13 min. Apical lengths and apical actin intensities of ablated and neighboring cells at different timepoints were stitched together to generate kymographs of the combined apical surface of the three cells (1 ablated + 2 neighbors).

Note that laser ablation did not disrupt the integrity of the cell membrane. No indication of a punctured membrane or leakage of cytoplasmic material was observed in brightfield images during laser ablation experiments for both smaller and larger epiblasts.

### Two-dimensional image analysis

#### Lumen solidity

Synthetic epiblasts were first incubated with fluorescent dextran (see previous section on tight junction permeability studies) to visualize smaller lumens. Outlines of lumen were manually drawn in ImageJ using fluorescent dextran or brightfield images. Lumen size and shape metrics including lumen solidity were measured in ImageJ. Lumen radius was determined from measured lumen area assuming a perfect circle (see equation [1]).

#### Nuclear area and perimeter

Nuclear area and perimeter of synthetic and human epiblasts as well as hiPSCs in 2D culture, were respectively measured from immunostained images acquired in-house and from previously published sources (Deglincerti et al., 2016; Molè et al., 2021; Nakagawa et al., 2014; Rodin et al., 2010; Shahbazi et al., 2016; Simunovic et al., 2019; Zhang et al., 2014). Using ImageJ, nuclear images were thresholded, smoothened using median filter (radius of 4 pixels), and processed using a Watershed algorithm (ImageJ) to separate touching nuclei. Nuclear area and perimeter were then measured in ImageJ.

#### Cell thickness of synthetic epiblasts

For quantifying average thickness of cell layer in smaller and larger epiblasts, 5 lines were manually drawn per cluster from the apical to the basal surface and their corresponding lengths were measured from immunostained images of whole synthetic epiblasts in ImageJ. These 5 lengths were averaged and reported as the average cell thickness.

### Three-dimensional image analysis

#### Lumen volume, number of cells, apical surface area and cell volume quantification

High resolution z-stacks of nucleus and actin in immunostained whole synthetic epiblasts were acquired (see previous section on immunofluorescence of whole synthetic epiblasts) using a Leica SP8 laser scanning confocal microscope with a HC FLUOTAR L 25×/0.95 NA water immersion objective. Images were acquired with a voxel size of 0.0909 × 0.0909 × 0.5691 µm^3^ (x y z). Lumen volume, number of cells, apical surface area of individual cells and individual cell volumes of synthetic epiblasts were quantified from these high-resolution z-stacks using Imaris 9.9 software (Bitplane). Nuclear images were used to quantify number of cells with Surfaces program in Imaris. Actin images were used to quantify lumen volume and apical surface area of individual cells with Surfaces program in Imaris. Nucleus and actin images were both used to segment individual cells and quantify cell volumes with Cells program in Imaris. Lumen radius was determined from measured lumen volume assuming lumen to be a perfect sphere.

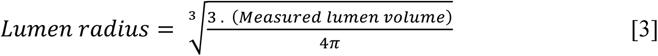

#### Lumen curvature quantification

To quantify lumen surface curvature in smaller and larger epiblasts, lumen surfaces reconstructed in Imaris from high-resolution z-stacks of immunostained whole synthetic epiblasts were exported to ImageJ. LimeSeg plugin in ImageJ was then used to compute gaussian curvature at each point of lumen surface.

### Morphological comparison of human and synthetic epiblasts

To compare morphological features of human and synthetic epiblasts, number of cells and lumen volumes of synthetic epiblasts on different days of culture were compared to those of human epiblasts. Average number of cells in human epiblasts on different days post fertilization were previously reported (Shahbazi et al., 2016). To quantify the evolution of lumen volumes as a function of number of cells in human epiblasts (Fig. 1E), previously published data and images were used. Lumen volumes were estimated from previously reported radius of gyration values of human epiblasts (Simunovic et al., 2019), assuming human epiblasts to be a perfect sphere and individual cell volume to be ~1500 µm^3^ (which was the average cell volume in synthetic epiblasts). For these human epiblasts, number of cells were reported (Simunovic et al., 2019). Next, to obtain additional data points from human epiblasts, we analyzed published 2D immunostained images of human epiblasts (Deglincerti et al., 2016; Molè et al., 2021; Shahbazi et al., 2016; Shahbazi et al., 2017; Simunovic et al., 2019) and measured lumen area and total cell area in the mid-plane of human epiblasts. This 2D lumen area and total cell area in mid-plane were used to estimate 3D lumen volume and number of cells in the human epiblast using the following assumptions: (i) human epiblast tissue and lumen are perfectly spherical, (ii) each cell in human epiblast has a cell volume of ~1500 µm^3^ (which was the average cell volume in synthetic epiblasts).

### Analysis of single cell RNA sequencing data

Human embryo derived single cell RNA sequencing datasets (Xiang et al., 2020; Zhou et al., 2019) were analyzed using Seurat R package (v4.2.0). Unprocessed datasets were obtained from publicly available GEO repositories: GSE136447 (Xiang et al., 2020) and GSE109555 (Zhou et al., 2019). Datasets were filtered to only include epiblast cells based on previously annotated populations in respective studies. Default setups in Seurat were used unless noted otherwise. Cells with ≤4,500 genes detected were discarded from analysis. Gene expression was calculated by normalizing raw counts by the total count, multiplying by 10,000 and performing log-transformation. Then, principal component analysis was performed in Seurat. Cell clusters were identified by a K-nearest neighbor (KNN) clustering approach with a resolution of 0.8. Non-linear dimensionality reduction was performed using Uniform Manifold Approximation and Projection (UMAP) algorithm (dimensions 1 to 10). UMAP showed that cell clusters obtained by KNN based clustering were roughly separated based on embryo age (days post fertilization) and were annotated accordingly. Finally, average expression level of different genes of interest were plotted for each cell cluster.

### Statistical analysis

Statistical analyses were performed using GraphPad Prism 9.3.1 software. Bar plots and respective error bars are defined throughout the figures. Statistical tests used, *n* and *P* values are listed in figure captions. *P* values less than 0.05 were considered statistically significant. All tests used were two-tailed unless mentioned otherwise.

In Fig. 1, synthetic epiblast immunostaining images (Fig. 1, B and C) are representative of 3 independent biological replicates. In Fig. 2, images (Fig. 2, C, F, G, H and K) are representative of 3 independent biological replicates. In Fig. 3, images (Fig. 3J) are representative of ≥3 independent biological replicates. In Fig. S1, synthetic epiblast immunostaining images (Fig. S1, B, C, F and G) are representative of 3 independent biological replicates. In Fig. S2, synthetic epiblast immunostaining images (Fig. S2H) are representative of 3 independent biological replicates. In Fig. S3, synthetic epiblast immunostaining images (Fig. S3A) are representative of 3 independent biological replicates. For all experimental plots, data are pooled from ≥3 biological replicates unless mentioned otherwise in the figure captions.

### Theoretical model and computational simulations

Description of the main hypotheses and equations of the physical models and the methods underlying the numerical simulations are provided in supplementary information.

## Supplementary Materials

- Figures S1 to S6
- Videos S1 to S9
- Supplementary Methods (procedures for computational modeling)

**Figure S1.**
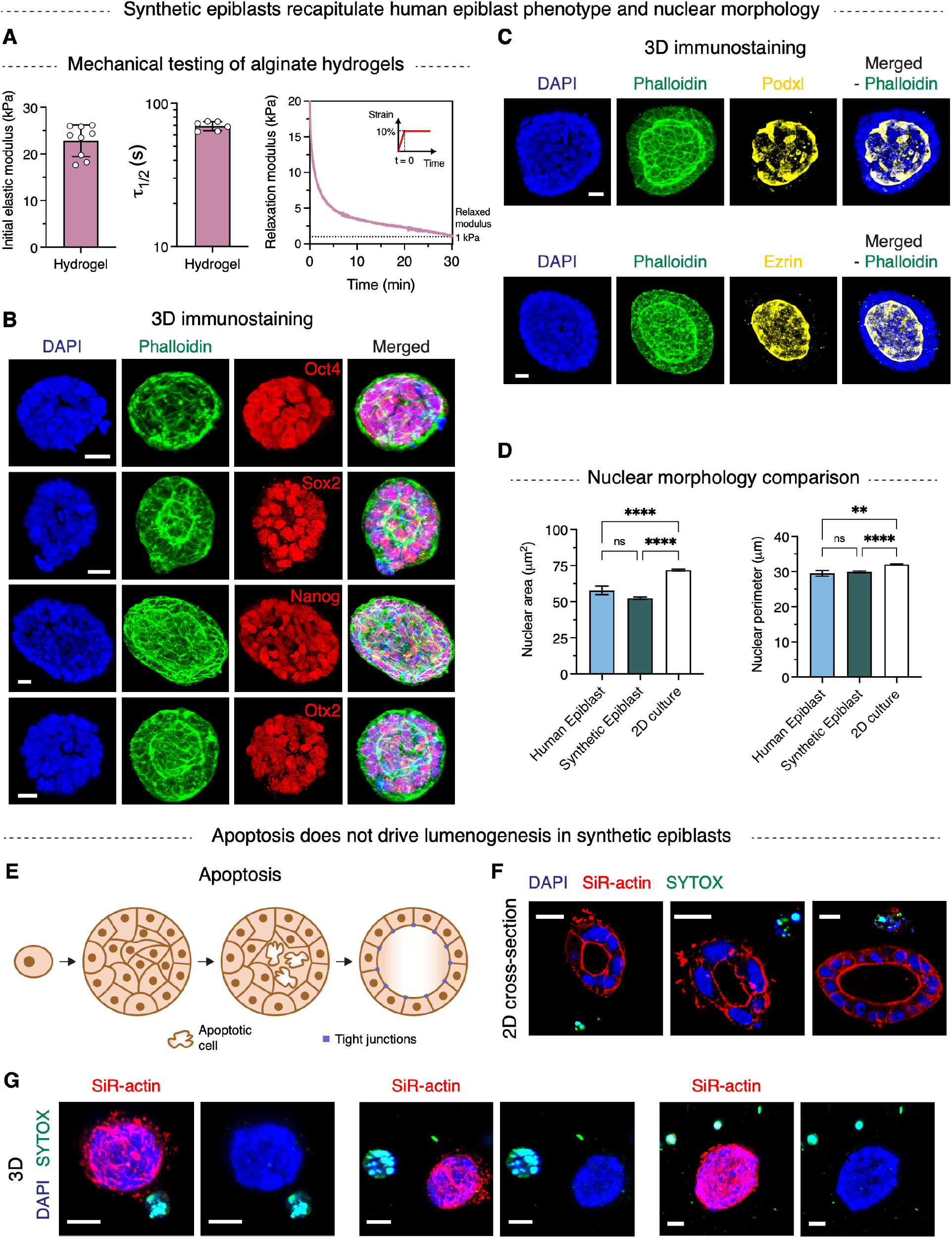
Synthetic epiblasts model human epiblasts and apoptosis does not drive lumenogenesis in synthetic epiblasts. **(A)** Mechanical characterization of viscoelastic alginate hydrogels using compression testing. Measurements of initial elastic modulus and stress relaxation were performed (mean ± s.d.; *n* ≥ 6). τ_1/2_ is the timescale of stress relaxation, defined as the time when the relaxation modulus reaches 50% of initial value. Representative stress relaxation profile of alginate hydrogels at 10% compressive strain is shown. Inset indicates applied strain profile. Final relaxed modulus of alginate hydrogels is ~1 kPa. **(B)** Maximum intensity projections of 3D immunostains of pluripotency markers Oct4, Sox2, Nanog (core pluripotency), Otx2 (primed pluripotency), phalloidin (actin) and DAPI (nucleus) in synthetic epiblasts. **(C)** Maximum intensity projections of 3D immunostains of Podocalyxin, Ezrin (apical polarity), phalloidin (actin) and DAPI (nucleus) in synthetic epiblasts. **(D)** Quantification of nuclear area and perimeter in human and synthetic epiblasts, as well as in 2D culture of hiPSCs (mean ± s.e.m.; ****p < 0.0001, **p < 0.01, ns: not significant p > 0.05, one-way ANOVA; *n* = 137 (human epiblast nuclei), 435 (synthetic epiblast nuclei), 2607 (2D culture nuclei)). See materials and methods section for sources of human epiblast and 2D culture images. **(E)** Schematic of apoptosis driven lumenogenesis. **(F)** SYTOX assay. Fluorescence images of live cells stained with SYTOX (dead cells), SiR-actin (actin) and DAPI (nucleus). **(G)** Maximum intensity projections of 3D stains of SYTOX (dead cells), SiR-actin (actin) and DAPI (nucleus). Scale bars: 20 μm (B, C, F, G).

**Figure S2.**
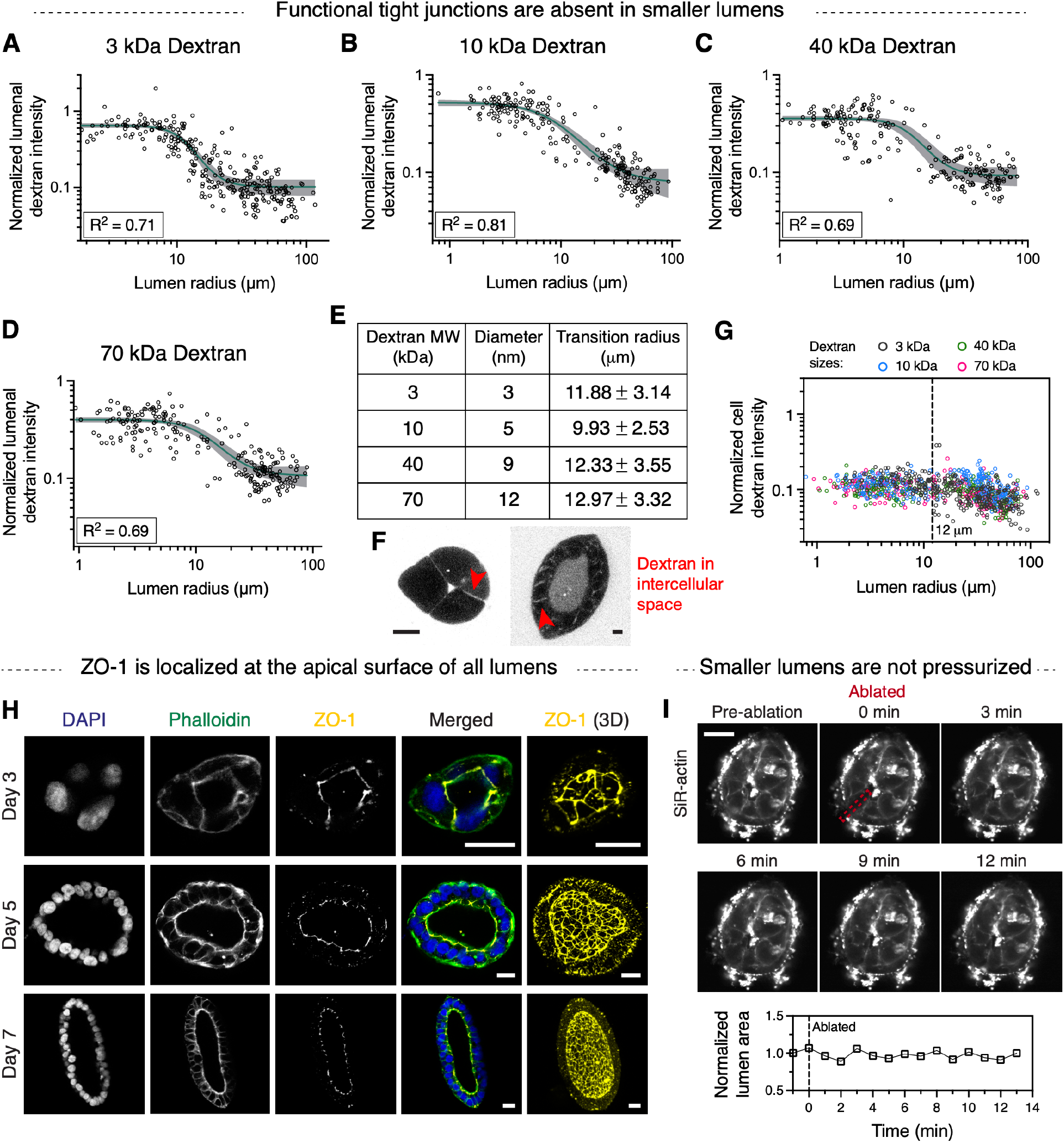
Functional tight junctions are absent in smaller epiblasts. **(A-D)** Individual plots showing quantification of dextran intensity inside lumen for different dextran sizes: 3 kDa (A), 10 kDa (B), 40 kDa (C), 70 kDa (D). Lines indicate sigmoidal fits and 95% CI. (*n*, R^2^) = 3 kDa: (290, 0.71), 10 kDa: (229, 0.81), 40 kDa: (213, 0.69), 70 kDa: (228, 0.69). **(E)** Table summarizing dextran sizes (from manufacturer) and respective IC50 or sigmoid fit inflection point values. **(F)** Representative fluorescence image of dextran (3 kDa) in smaller lumens. Image brightness was increased to visualize dextran in the intercellular spaces. **(G)** Quantification of fluorescent dextran intensity inside cells normalized to that in the hydrogel. This intensity value represents noise in imaging, as dextran is cell impermeable. **(H)** Immunostains and maximum intensity projections of 3D immunostains of ZO-1 (tight junction protein), phalloidin (actin) and DAPI (nucleus) in synthetic epiblasts on different days of culture. **(I)** Laser ablation through an entire cell and quantification of lumen area pre- and post-ablation. No large change in lumen area is observed post-ablation, suggesting that smaller lumens are not pressurized. Scale bar: 10 μm (F), 20 μm (H, I).

**Figure S3.**
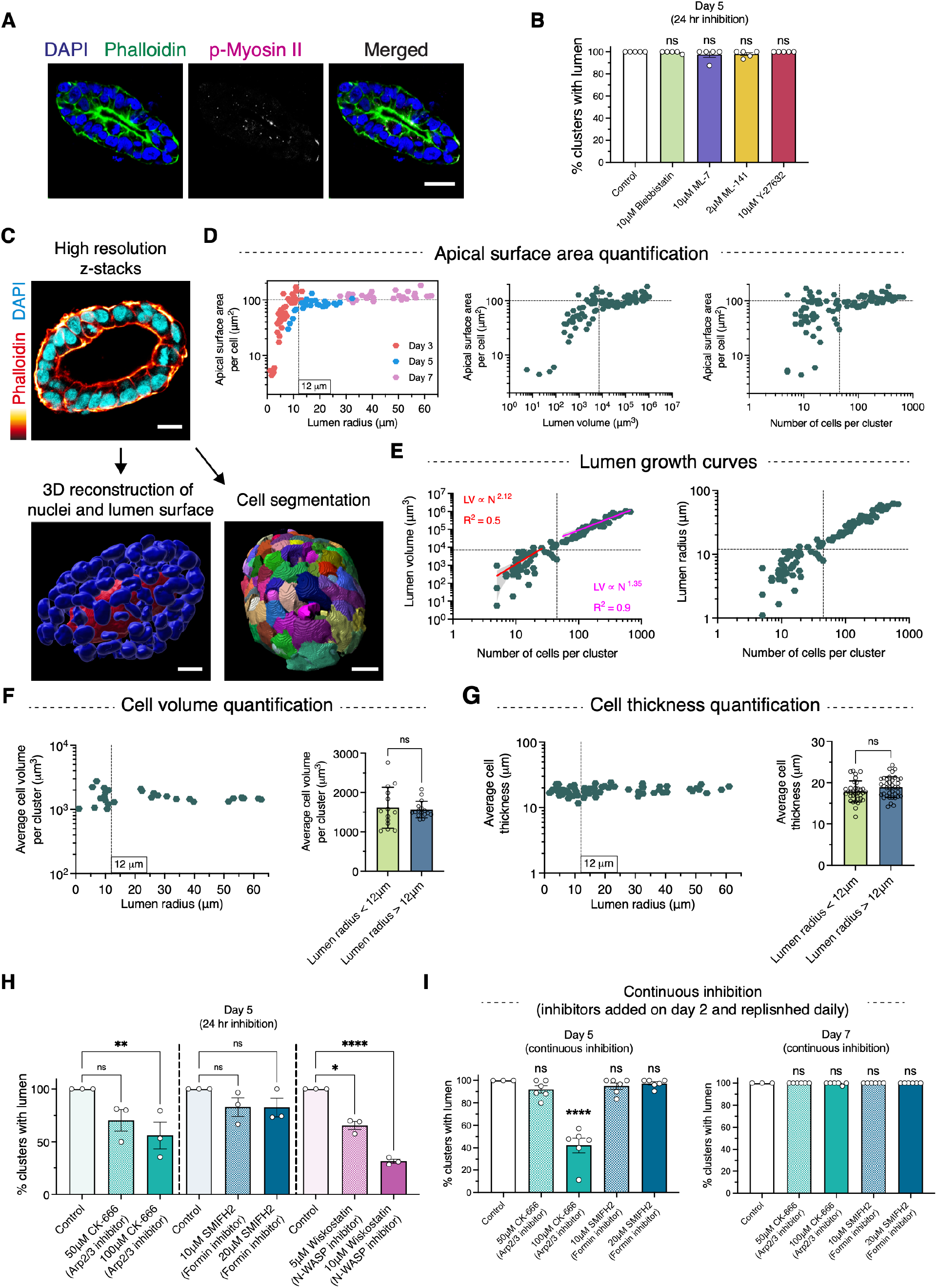
Actin polymerization is necessary for lumenogenesis and the two size-dependent mechanisms of lumen growth show distinct growth dynamics. **(A)** Immunostains of phosphorylated myosin II, phalloidin (actin) and DAPI (nucleus) in synthetic epiblasts. **(B)** Percent epiblasts with lumen in the presence of different actomyosin contractility inhibitors on day 5 of culture (mean ± s.e.m.; ns: not significant p > 0.05, one-way ANOVA; *n* = 5). **(C)** Image analysis pipeline for quantification of lumen and cell size metrics from 3D z-stacks of phalloidin (actin) and DAPI (nucleus) stained synthetic epiblasts. **(D)** Quantification of average apical surface area per cell as a function of lumen radius, lumen volume or number of cells (*n* = 101 synthetic epiblasts). **(E)** Quantification of lumen growth dynamics (*n* = 101 synthetic epiblasts). Smaller lumens grow faster than large lumens for a fixed number of cells (N). Lines indicate power law fits. For smaller lumens, lumen volume (LV) μ N^2.12^ (R^2^ = 0.5) and for larger lumens, LV μ N^1.35^ (R^2^ = 0.9). **(F)** Quantification of cell volume as a function of lumen radius (*n* = 30 synthetic epiblasts). **(G)** Quantification of cell layer thickness as a function of lumen radius (*n* = 71 synthetic epiblasts). **(H-I)** Percent epiblasts with lumen in the presence of Arp2/3 (CK-666), formin (SMIFH2) and N-WASP (Wiskostatin) inhibitors for either 24 hr (H) or continuously from day 2 onwards (I) (mean ± s.e.m.; ****p < 0.0001, **p < 0.01, *p < 0.05, ns: not significant p > 0.05, one-way ANOVA; *n* = 3). Scale bar: 20 μm (A, C).

**Figure S4.**
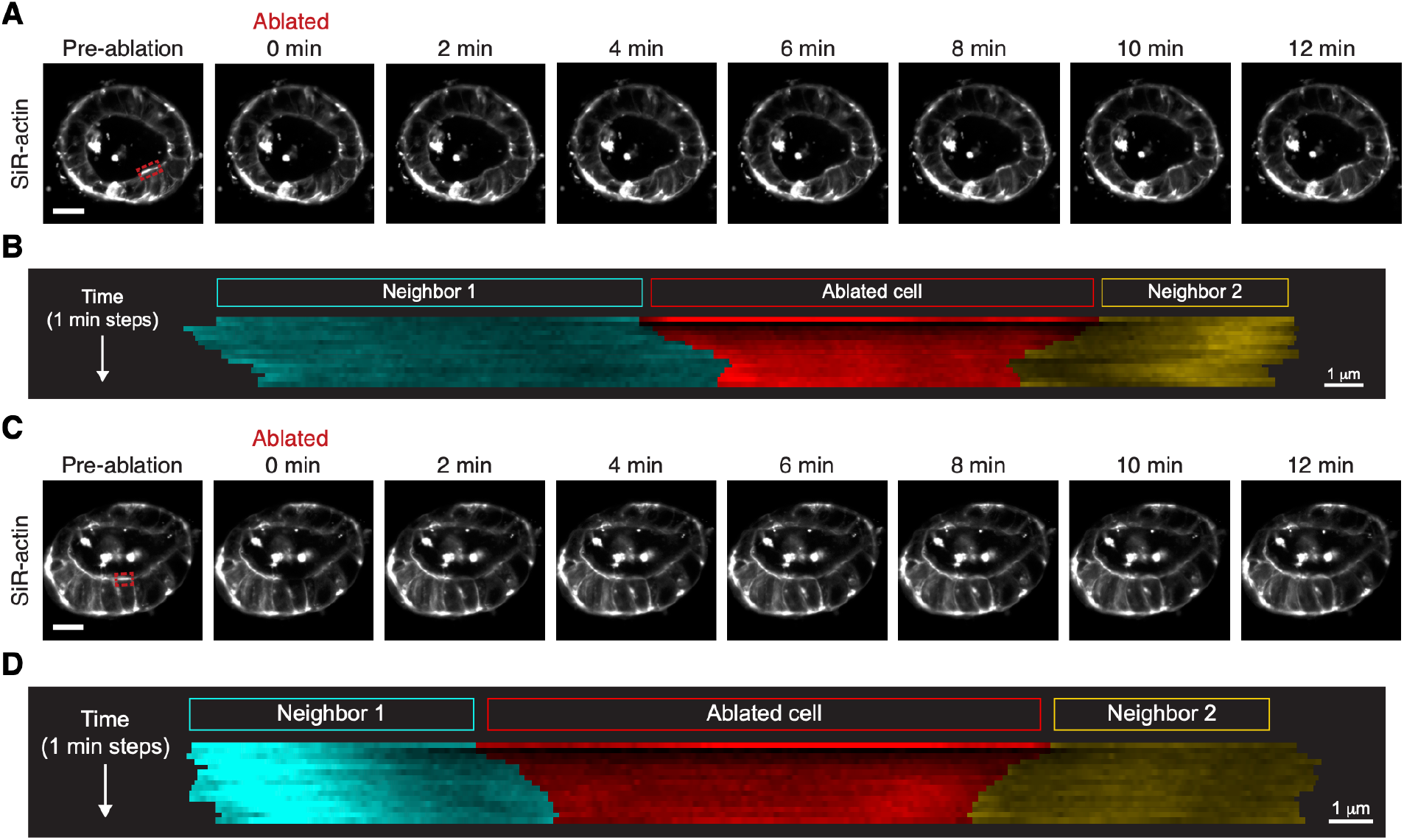
Apical actin ablation in smaller epiblasts results in apical length shrinkage in the ablated cell and expansion in the neighboring cells. **(A, C)** Apical actin ablation and recovery. Red outline indicates ablated surface. **(B, D)** Kymograph showing apical actin of ablated and neighboring cells for epiblasts shown in (A and C) respectively. Scale bars: 20 μm (A, C), 1 μm (B, D).

**Figure S5.**
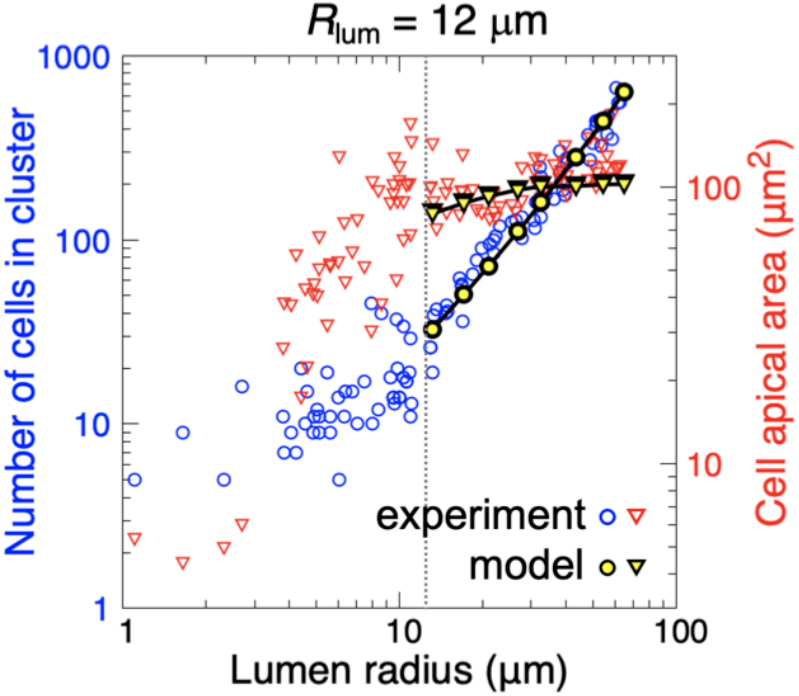
Physical model of osmotic pressure driven lumen growth in larger epiblasts. Model predictions closely match experimental observations of lumen radius, number of cells and cell apical surface area of larger epiblasts.

**Figure S6.**
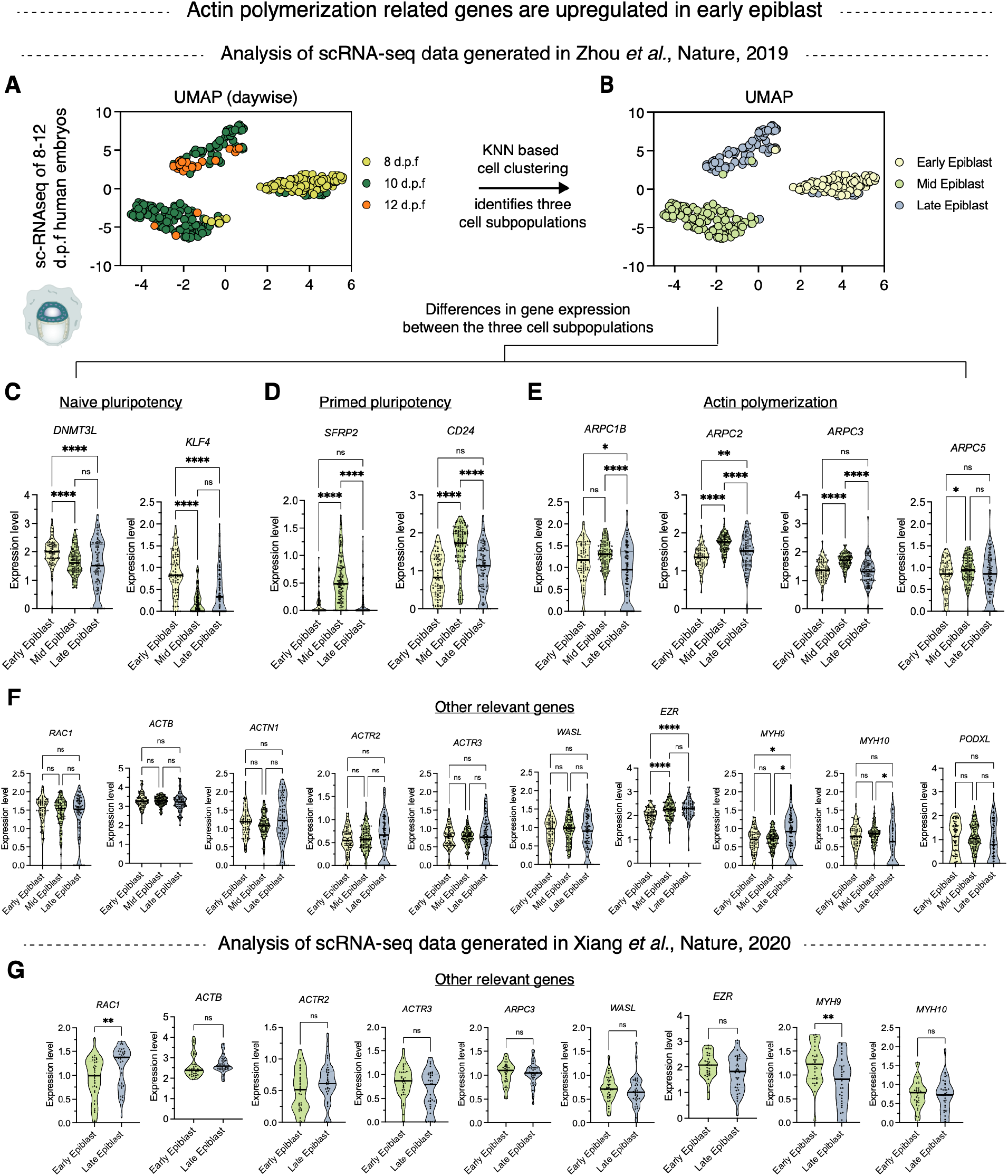
Human epiblasts show upregulation of actin polymerization related genes during 8-10 d.p.f. **(A)** UMAP plot of human epiblast cells generated from scRNA-seq data of peri-implantation human embryos (Zhou et al., 2019) (274 epiblast cells) colored with timepoints (d.p.f) **(B)** KNN (k-nearest neighbor) based cell clustering. The three subpopulations are annotated as early, mid and late epiblast. **(C-F)** Normalized expression level of naïve pluripotency genes (C), primed pluripotency genes (D), actin polymerization related genes (E) and other relevant genes (F) (median and quartiles; ****p < 0.0001, **p < 0.01, *p < 0.05, ns: not significant p > 0.05, Kruskal-Wallis; *n* = 78 (early epiblast cells), 111 (mid epiblast cells), 85 (late epiblast cells)). **(G)** Normalized expression level of genes associated with actin polymerization and actomyosin contractility for the scRNA-seq data (Xiang et al., 2020) shown in Fig. 6, A to E (median and quartiles; **p < 0.01, ns: not significant p > 0.05, Mann-Whitney; *n* = 34 (early epiblast cells), 37 (late epiblast cells)).

**Video S1. z-stack of synthetic epiblasts on day 7 of culture, immunostained for Oct4, ZO-1, actin (phalloidin), and nucleus (DAPI).**

**Video S2. Fluorescence recovery after photobleaching (FRAP) of lumenal dextran.** Red arrowhead indicates bleached lumenal dextran.

**Video S3. Brightfield timelapse imaging of lumen opening and growth in a smaller epiblast.** Red arrowhead indicates lumen opening and subsequent growth.

**Video S4. Brightfield timelapse imaging of lumen growth in a larger epiblast.**

**Video S5. Super-resolution imaging z-stack of actin in a day 6 synthetic epiblast.**

**Video S6. Actin and brightfield timelapse imaging of lumen growth in a smaller epiblast.**

**Video S7. Apical actin ablation and recovery in a smaller epiblast.** Dotted red box indicates ablated apical surface. Red arrowhead follows the ablated cell during the timelapse. Cyan arrowhead indicates the absence of blebbing post-ablation.

**Video S8. Actin and brightfield timelapse imaging of lumen growth in a larger epiblast.**

**Video S9. Apical actin ablation and recovery in a larger epiblast.** Dotted red box indicates ablated apical surface. Red arrowhead follows the ablated cell during the timelapse. Magenta arrowhead indicates the formation of a bleb post-ablation.

