## Supplementary Methods for "Actin polymerization drives lumen formation in a human epiblast model"

This file contains:

- Supplementary Methods (procedures for computational modeling)

### Supplementary Methods

In this Supplementary Methods section, we introduce the main hypotheses and equations of the physical models and the methods underlying the numerical simulations.

### Model of lumen growth

As experiments indicated two lumen-size dependent mechanisms of lumen growth (Supplementary Methods Figure 1), we modeled each of the mechanisms to better understand the physics involved. Smaller epiblasts do not have functional tight junctions and thus clefts between cells are highly penetrable to ions and water. However, there is still spontaneous opening of lumens associated with extensive actin polymerization on the apical side of cells enclosing lumen (Supplementary Methods Figure 1). Until formation of tight junctions in larger epiblasts, there is no significant accumulation of ions in the lumen even at large ion pumping due to free diffusion of ions through the clefts. In larger epiblasts there is no apical actin polymerization and formation of tight junctions enables accumulation of ions in the lumen. Hence, we built a model of lumen growth due to ion pumping and osmotic pressure buildup that is representative of the mechanism responsible for growth of larger lumens. We find that this model accurately captures observed lumen growth in larger epiblasts. However, when sufficient leakage of ions along the intercellular spaces was present, as was the case in smaller epiblasts, osmotic pressure did not buildup and no lumen growth was observed. As these epiblasts have apical actin polymerization which can generate stress to open and grow lumens, we added apical actin polymerization to our model which resulted in lumen opening and growth and predicted features of the apical actin mesh that are necessary for lumen growth.

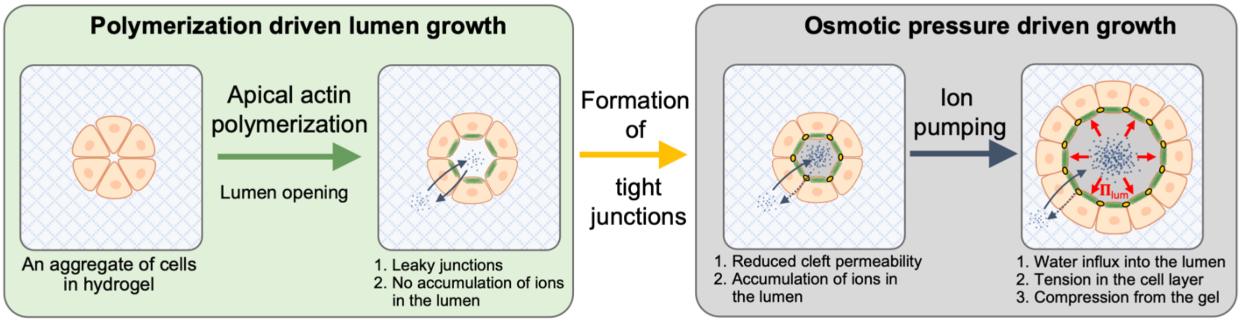

**Supplementary Methods Figure 1: Schematic of the two distinct lumen-sized dependent mechanisms of lumen growth.**

### A. Analytic predictions

For the purpose of analytical treatment, we consider a simplified geometry of a lumen-containing synthetic epiblast. The monolayer of cells surrounding a lumen is represented by a collection of spheres of radius $R_{cell}$ that in perfectly symmetrical lumens are tightly packed on a spherical surface of a larger radius. Due to the rotational symmetry, we can first consider the equatorial cross-section of the synthetic epiblast consisting of $n_{cells}$ spheres (Supplementary Methods Figure 2A). To account for the cell-cell adhesion, the spheres are allowed to partly overlap forming a contact line of length $L_{cleft}$, which represents the effective surface tension in cells and depends on the adhesion strength. Larger adhesion between cells would result in larger contact length and higher contact angle. If contact between cells $L_{cleft}$ remains constant, then the lumen radius, $R_{lum}$, and the apical length, $L_{apical}$ can be defined as

$$R_{lum} = cot\frac{\pi}{n_{cells}}r - \frac{L_{cleft}}{2} \quad [1]$$

$$L_{apical} = R_{cell}(\pi - \frac{2\pi}{n_{cells}} - 2\arccos(\frac{r}{R_{cell}})) \quad [2]$$

where $r = \sqrt{R_{cell}^2 - L_{cleft}^2/4}$ is the distance between the cell center and the cleft.

The dependence of the normalized lumen radius on the apical length of cells at different cleft lengths are shown in Supplementary Methods Figure 2B. Assuming close packing of cells on a spherical surface, approximately each cell has an apical surface area $A_{apical} \approx 2\sqrt{3}\, R_{cell}^2\, sin^2(\frac{L_{apical}}{2\, R_{cell}})$ at radius $R_{lum}$ from the sphere center. When cleft length is decreased (corresponds to reduced adhesion between cells), the synthetic epiblast is predicted to demonstrate spontaneous lumen opening due to increasing apical length. The apical growth rate is shown to be much higher in smaller epiblasts (Supplementary Methods Figure 2C) reaching the maximum in lumens of size ~3 μm and monotonically decreasing as lumen grows. This spontaneous lumen opening is associated with only a small increase in the number of cells, where the predicted number of cells in a spherical epiblast is given by $N_{cells} \approx 4\pi R_{lum}^2 / A_{apical}$. The model prediction is in a good agreement with experiment (Supplementary Methods Figure 2D) and shows that rapidly increasing apical length causes lumen expansion at earlier stages of lumen growth in synthetic epiblasts.

Since the synthetic epiblast is surrounded by a gel that creates a confinement for the growing epiblast, we estimate the critical lumen size where the epiblast becomes unstable due to the pressure from the gel, at which point it will tend to buckle inwards in order to reduce the gel deformation. The gel pressure is proportional to the Young's modulus $E_{gel}$ of the gel and the amount of strain in it $\varepsilon_{gel}$, and reads as

$$P_{gel} = E_{gel}\varepsilon_{gel} \approx E_{gel}(\frac{R_{out}}{R_{out}^{ini}} - 1) \quad [3]$$

where $R_{out} = \frac{R+R_{cell}+R_{lum}+L_{cleft}}{2}$ is the average epiblast radius on the outer side of cells. An approximation for the critical buckling pressure for an epithelium monolayer of thickness $h$, Young's modulus $E$, and Poisson ration $\nu$ can be written as

$$P_{buckling} \approx \lambda^{\frac{2}{5}} K^{\frac{3}{5}} / R^{\frac{11}{5}} \quad [4]$$

where $\lambda = Eh/(1-\nu^2)$, $K = Eh^3/(1-\nu^2)$ are compressional and bending rigidities of the monolayer, respectively, and $R$ is the radius (Trushko et al., 2020). For a cell monolayer of thickness $L_{cleft} = R_{cell} = 6$ µm, $E = 20$ kPa and a gel with modulus $E_{gel} = 1$ kPa, a buckling transition is predicted to occur at $R_{lum}/R_{cell} \approx 1.95$, which corresponds to $R_{lum} \approx 12$ µm. This means that large epiblasts are required to build higher pressure in the lumen to stabilize the shape, but when compression from the gel is small enough, lumen opening does not require pressurized lumens.

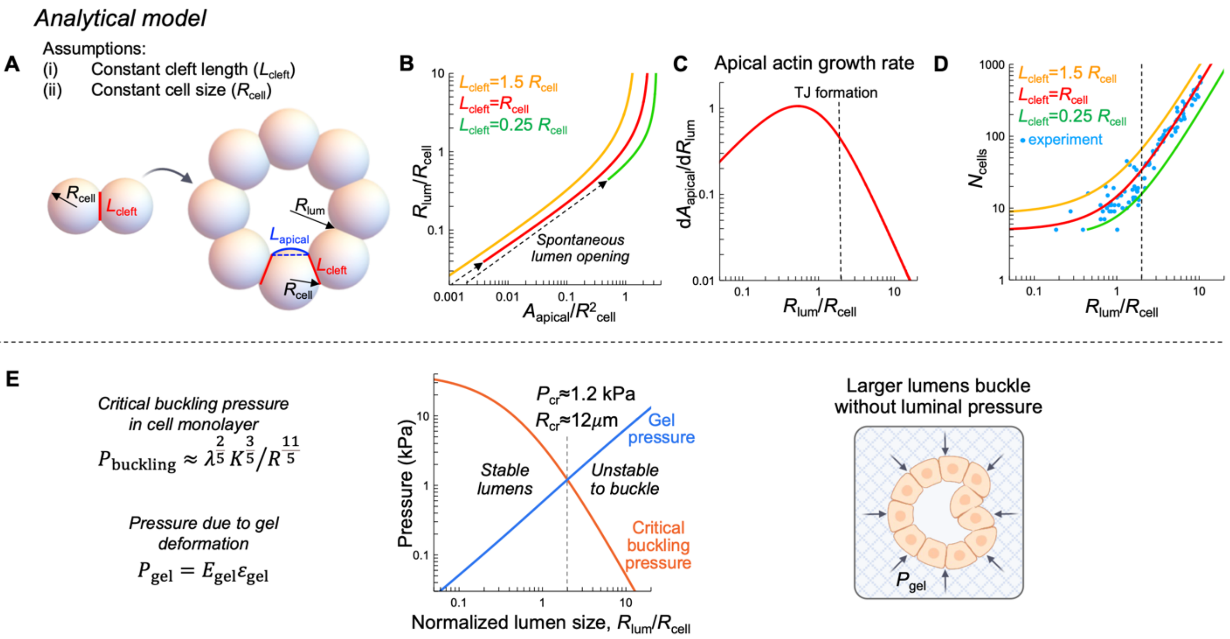


**Supplementary Methods Figure 2: Analytical model. (A-D)** Schematic of a simple analytical model for estimating evolution of cell apical surface area and cell number with increase in lumen size, for fixed cell volume and cell thickness or cleft length. **(E)** Buckling pressure considerations predict a transition to pressure-driven lumen growth at ~12 μm lumen radius to prevent cell layer buckling.

#### *Apical actin polymerization generates stresses in the cortical layer to counteract gel confinement and drive lumen growth*

As smaller epiblasts lack tight junctions that prevents the buildup of osmotic pressure, we considered the effect of actin polymerization on lumen growth (Supplementary Methods Figure 3A, B) To elucidate the effect of actin polymerization, we calculate the hydrostatic pressure difference between the lumen and the external media. Lumen opening is associated with water flux into the lumen, which in the absence of osmotic gradients is possible if hydrostatic pressure in the lumen, $P_{lum}$ is smaller than the external hydrostatic pressure, $P_{ext}$. Then, lumen growth will be proportional to the difference in pressure

$$P_{lum} - P_{ext} = \Delta P_b - \Delta P_a \quad [5]$$

where $\Delta P_b, \Delta P_a$ are the gradients in the hydrostatic pressure at basal and apical side of the cell, respectively. Using the Laplace law, we calculate the gradients as

$$\Delta P_b = \frac{2h_b(\sigma_b^p + \sigma_b^a)}{R_{cell}} + P_{gel} \quad [6]$$

$$\Delta P_a = \frac{2\xi_a \widetilde{h_a}(\sigma_a^p + \sigma_a^a)}{R_{cell}} \quad [7]$$

$$\sigma_a^p = K_a(\frac{L_a}{\xi_a \widetilde{L_a}} - 1) \quad [8]$$

$$\sigma_b^p = K(\frac{L_b}{\widetilde{L_b}} - 1) \quad [9]$$

where $\sigma_a^p, \sigma_b^p$ are passive stresses in the cortex, and $\sigma_a^a, \sigma_b^a$ are active stresses due to actomyosin contractility that are assumed to be constant, $h_b, h_a$ are thicknesses of the cortical layer, and $K_a, K$ are effective stiffnesses of the cortex on the respective sides of the cell (Supplementary Methods Figure 3A). The passive stress on the apical side depends also on the undeformed length factor, $\xi_a$, which changes the reference (undeformed) size of the cortex to account for its growth without deformation and thus without stress generation (Supplementary Methods Figure 3A). During polymerization the cortex not only elongates but can also stiffen due to crosslinking between filaments in the network. In Supplementary Methods Figure 3C we plot the dependence of the lumen growth rate on both cortex stiffness ($K_a$) and undeformed length factor ($\xi_a$) assuming constant strain in the cortex ($\frac{L_b}{\widetilde{L_b}} = 1, \frac{L_a}{\widetilde{L_a}} = 12$). Increasing undeformed length factor ($\xi_a$) leads to decreased tension on the apical side and, as a result, the pressure difference also decreases, whereas increasing stiffness ($K_a$) leads to the opposite effect and lumen expansion is predicted to be faster. Since it is experimentally challenging to measure the actual stiffness and the undeformed length factor in the cortex, we expect that they are both increased but stiffness increase outcompetes any increase in undeformed length factor to drive lumen growth.

Overall, the above model predictions allow us to conclude that (i) apical actin polymerization drives lumen growth in smaller epiblasts, (ii) cell size and cell contact or cleft length remain constant during lumen growth, which we verified experimentally, and (iii) in larger epiblasts, lumen pressure is required to balance the pressure from the gel to avoid buckling. To examine the lumen growth at later stages, where lumen pressure is necessary to avoid buckling, we employ a model for pressure-driven lumen growth that accounts for ion and water fluxes through the cell layer and the intercellular space and results in pressure gradients sufficient to drive lumen growth.

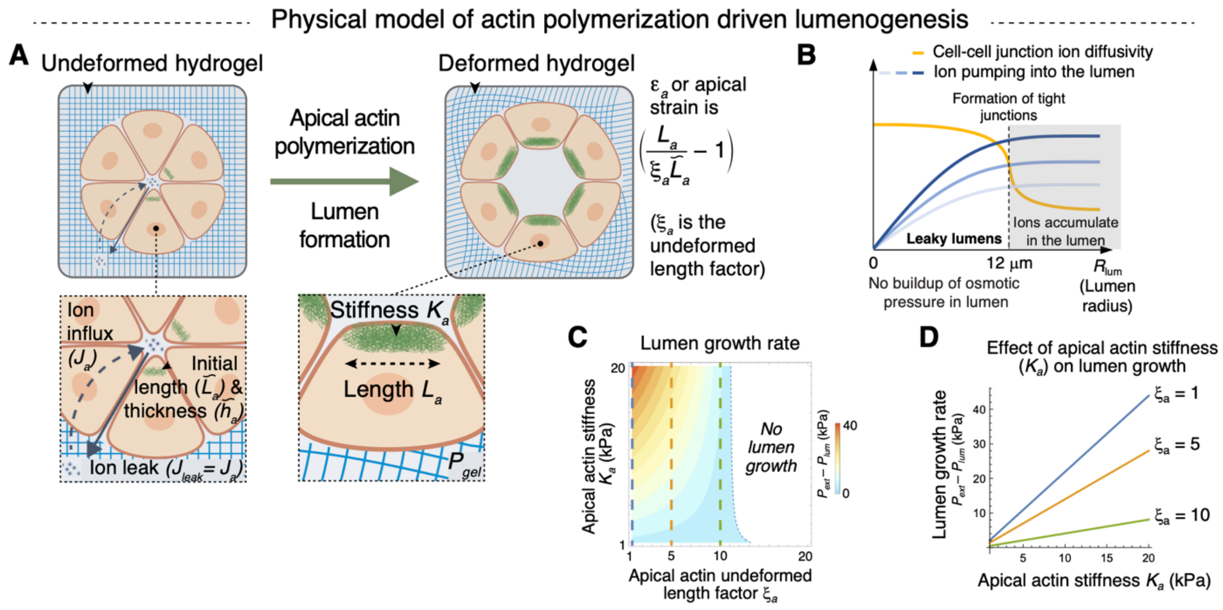


**Supplementary Methods Figure 3: Physical model of lumenogenesis in small synthetic epiblasts. (A)** Schematic of the theoretical model of actin polymerization driven lumen growth. $\tilde{L}_a$: initial length of apical actin mesh; $L_a$: final length of apical actin mesh; $\xi_a$: undeformed length factor; $K_a$: stiffness of apical actin mesh; $J_a$: ion flux into lumen; $J_{leak}$: ion leak along the leaky junctions; $P_{gel}$: stress exerted by the hydrogel on the cell cluster. **(B)** Model shows that osmotic pressure does not build in leaky lumens. Ions pumped into the lumen diffuse out along the leaky junctions in smaller epiblasts preventing buildup of osmotic pressure. **(C, D)** Plots showing that increase in apical actin stiffness ($K_a$) and decrease in undeformed length factor ($\xi_a$) result in higher lumen growth rates. Lumen growth rates as a function of apical actin stiffness ($K_a$) for given values of undeformed length factor ($\xi_a$) are shown in (C). All other parameters were kept constant. $h_a = h_b = 0.6$ µm, $R_{cell} = 6$ µm, $L_a/\tilde{L}_a = 12$, $L_b/\tilde{L}_b = 1$.

### B. Pressure driven lumen growth in larger synthetic epiblasts

Since formation of tight junctions between cells is observed only in synthetic epiblasts larger than 12 µm, the permeability of clefts to ions and water is large at early stages, and, thus, ions do not accumulate in the lumen. In larger epiblasts with functioning tight junctions, the ion concentration in the lumen increases and becomes higher than in the external media. This builds up a concentration gradient across the cell layer and increases the osmotic pressure in the lumen that causes a flux of water into the lumen resulting in lumen expansion (Supplementary Methods

Figure 4A). To examine the lumen size in larger epiblasts that demonstrate growth even when there is an increasing pressure from the surrounding hydrogel as the lumen size increases, we include passive transport of ions and water as well as active ion pumping into the model (Supplementary Methods Figure 4B). We show that the transport through the cell membrane and clefts is regulated and in turn depends on both the lumen and cell size.

***Ion transport in the lumen and cells***

The change in the total number of ions in the cell is proportional to the total ion flux across the cellular membrane. The ion transport includes passive fluxes due to the difference in the osmotic pressure between the two sides of cell membrane, and the active fluxes that are provided through selective ion transport by ion pumps. The equilibrium osmotic pressure, $\Pi$, is related to the molar concentration, $c$, and the number of ions, $N^i$, by van't Hoff's equation as $\Pi = RTc = RTN^i/V$, where $R$ is the gas constant, $T$ is the temperature, and $V$ is the volume. Due to the apical-basal polarity in cells, the active fluxes are directional, and, in general, they are different at the basal and apical sides. The change in the number of ions in the cell is given by

$$d_t N_{cell}^i = J_b^i - J_a^i - 2J_c^i \qquad [10]$$

where $J_b^i = L_b(\omega \Delta\Pi_b + j_b)$, $J_a^i = L_a(\omega \Delta\Pi_a + j_a)$, $J_c^i = L_c \omega \Delta\Pi_c$ are the fluxes through the basal, apical and cleft side, respectively. The osmotic pressure differences, $\Delta\Pi_b = \Pi_{ext} - \Pi_{cell}$, $\Delta\Pi_a = \Pi_{cell} - \Pi_{lum}$, $\Delta\Pi_c = \Pi_{cell} - \Pi_{cleft}$ determine the passive ion transport through the cell membrane with ion permeability ω. The osmotic pressure inside the cleft can be averaged as $\Pi_{cleft} = (\Pi_{lum} + \Pi_{ext})/2$. The density of ion pumps and pumping rate, $j_a$, $j_b$ on both sides are assumed to be constant. Then total flux through a particular side of the cell is proportional to the respective lengths, $L_b, L_a, L_c$ for the basal, apical, and cleft sides.

The change in the total number of ions, $N_{lum}^i$ in the lumen is determined by the difference between influx of ions through the apical side of the cell and the leakage of ions through the clefts:

$$d_t N_{lum}^i = J_a^i - J_{leak}^i \qquad [11]$$

where the ion leak through the cleft is assumed to be proportional to the difference in concentrations between the lumen and the external solution, and it also depends on the ion diffusivity $D$ and the cleft width $h_c$ as $J_{leak}^i = \frac{D\, h_c}{L_c}(c_{lum} - c_{ext})$.

***Lumen and cell size regulation***

Due to the water incompressibility, the cell and lumen sizes change proportional to the total amount of water fluxes. The change in the cell size is then given by fluxes through the cell membrane

$$d_t A_{cell} = J_b^w - J_a^w - 2J_c^w \qquad [12]$$

The water fluxes are passive, and they are defined by hydrostatic, $\Delta P$, and osmotic pressure, $\Delta \Pi$, differences across the corresponding side of the cell membrane. Assuming that water permeability through the cell membrane, $\Lambda_m$, is constant, the fluxes through the basal, apical and cleft sides are $J_b^w = L_b \Lambda_m (\Delta \mathrm{P}_b - \Delta \Pi_b)$, $J_a^w = L_a \Lambda_m (\Delta \mathrm{P}_a - \Delta \Pi_a)$, and $J_c^w = L_c \Lambda_m (\Delta \mathrm{P}_c - \Delta \Pi_c)$, where $\Delta P_b = P_{ext} - P_{cell}$, $\Delta P_a = P_{cell} - P_{lum}$, and $\Delta P_c = P_{cell} - P_{cleft}$.

The change in the lumen size is proportional to the sum of water fluxes through the apical side and the leakage through the cleft

$$d_t A_{lum} = J_a^w - J_{leak}^w \qquad [13]$$

where $J_{leak}^w = \frac{\Lambda_{leak} h_c}{L_c}(\Delta P_{leak} - \Delta \Pi_{leak})$, $\Delta P_{leak} = P_{lum} - P_{ext} = -\Delta P_b - \Delta P_a$ and $\Delta \Pi_{leak} = \Pi_{lum} - \Pi_{ext} = -\Delta \Pi_b - \Delta \Pi_a$. The water leak through the cleft depends on permeability of the intercellular gap $\Lambda_{leak}$ to water and it can be different from the cell membrane permeability $\Lambda_m$.

***Stresses in the cortical layer and in the gel depend on pressure differences and determine the lumen size***

The hydrostatic pressure difference between the cell and external medium is determined by the Laplace law that relates the tension along the cell surface $\gamma$, and the curvature radius of the cell R with the pressure difference as $\Delta P = \frac{2\gamma}{R}$. Since the plasma membrane is a fluid-like structure, we assume that only stresses in the comparatively stiff cell cortex contribute to the tension $\gamma$.

The cortical stress on the apical side has passive, $\sigma_a^p$, and active, $\sigma_a^a$, components and can be written as

$$\sigma_a = \sigma_a^p + \sigma_a^a = \frac{R_a}{2h} \Delta P_a \qquad [14]$$

where $R_a$ is the curvature radius of the side, and $h$ is the cortex thickness. The active stress arises due to the actomyosin-mediated contractility in the cortex, and it is assumed to be constant.

The passive stress is caused by external load and for simplicity it can be assumed to be linearly proportional to the strain on the apical side,

$$\sigma_a^p = \frac{K_a}{2}\left(\frac{L_a}{\zeta_a L_a^{ini}} - 1\right) \quad [15]$$

where $K_a$ is the effective stiffness of the cortical layer, $L_a$, $L_a^{ini}$ are the actual and the initial lengths, respectively. The passive stress also depends on the polymerization on the apical side, which increases the reference length of this side by factor $\zeta_a \geq 1$.

The stress on the basal side can be written as

$$\sigma_b = \sigma_b^p + \sigma_b^a = -\frac{R_b}{2h}(\Delta P_b + \sigma_{gel}) \quad [16]$$

where $\sigma_b^a$ is the constant active stress, $\sigma_b^p = \frac{K}{2}\left(\frac{L_b}{L_b^{ini}} - 1\right)$ is the passive stress, and the compressive stress from the gel, $\sigma_g$ , on the basal side acts in addition to the external pressure and increases the hydrostatic pressure in the cell. The normal stress on the basal side due to deformations in the gel with effective stiffness $E_g$ is proportional to the increase in the lumen size and given by

$$\sigma_{gel} = E_{gel}\varepsilon_{gel} = \frac{E_{gel}}{2}\left(\frac{R_b}{R_b^{ini}} - 1\right) \quad [17]$$

This acts to decrease the surface tension caused by the pressure difference across the basal side and builds up a higher hydrostatic pressure in the cell. In the initial state, when there are no passive stresses ( $\sigma_b^p = 0$, $\sigma_{gel} = 0$ ), there still has to be a small unavoidable hydrostatic pressure difference $\Delta P_b = -\frac{2h\sigma_b^a}{R_b^{ini}}$ required to balance the constant active stress produced by myosin motors. We note that the assumption of the material as linearly elastic is a simplification of the viscoelastic and viscoplastic gels used in the experimental studies. We expect that our fundamental findings from the models would still hold, though the stresses remaining in the gel following lumen opening and synthetic epiblast expansion would be expected to relax in the experimental condition.

The cell shape and the number of cells ($n_{cells}$) in a perfectly symmetrical epiblast is defined by the sector angle $\phi = 2\pi/n_{cells}$. In larger synthetic epiblasts with increase in number of cells, each individual cell occupies a smaller sector but the lumen size increases. The curvature of the cell membrane on the basal side ($1/R_b$) depends on the apparent radius of the epiblast, whereas the apical curvature is assumed to be small, similar to the experimental observations.

Thus, the hydrostatic differences $\Delta P_b$ and $\Delta P_a$ depend both on the stresses in the cortex and in the gel that are defined by the lumen and cell sizes $A_{lum}$, $A_{cell}$, the number of cells n, and the curvature radius $R_a$ at the apical side.

***Simulation results for larger lumens***

Since the growth in larger lumens is predominantly caused by increasing osmotic pressure in the lumen rather than apical actin polymerization which becomes stabilized and does not provide enough mechanical force to expand lumens, we first examined the role of active ion pumping on the lumen size.

The mechanical equilibrium lumen size at constant number of cells depends both on apical and basal ion pumping (Supplementary Methods Figure 4C). For an epiblast consisting of 8 cells, the model predicts that the lumen forms only if there is a sufficient amount of ion influx into the cell through the basal side (>$0.3\times10^{-6}$ M/m$^2$/s). This builds up the necessary osmotic pressure in the cell, which maintains the cell size, cleft length and also increases the passive ion transport into the lumen. For basal ion pumping slightly above the minimal threshold (>$0.3\times10^{-6}$ M/m$^2$/s), small lumens can exist even without active pumping through the apical side. However, higher levels of basal ion pumping at small apical pumping restricts the lumen formation (purple region in Supplementary Methods Figure 4C) due to the large ion concentration in the cell that significantly increases accumulation of water in the cell and leads to increased cell size and higher pressure from the hydrogel. In addition, the osmotic pressure difference on the apical side becomes much larger than the hydrostatic pressure difference, leading to transport of water from the lumen into the cell and preventing lumen formation.

In the other limit, when ion pumping on apical side is greater than on the basal side, the model predicts a reduction in cell size due to decreased osmotic pressure in the cells. As a result, the cleft length decreases and ions do not accumulate in the lumen, preventing lumen formation (green region in Supplementary Methods Figure 4C).

In the intermediate regime at high enough basal ion pumping that is balanced by pumping ions out of the cell through the apical side, the lumen size is shown to be strongly dependent on the ratio between active transport on both sides. Interestingly, lumens are predicted to become larger with increasing pumping from the basal side because it increases both the passive ion flux into the lumen and the cleft. But the dependence of lumen size on pumping from the apical side changes based on basal ion pumping. When basal ion pumping is small, lumens increase in size with increase in apical pumping, but at higher basal pumping, lumens become smaller with increase in apical pumping.

In order to capture the experimental observations in large synthetic epiblasts, we simulated lumen growth in a system with large number of cells. For that, we fixed the number of cells and allowed the system to grow to an equilibrium size while keeping the rest apical and basal lengths constant (the rest lengths in undeformed state correspond to the case with 8 cells in the cluster). In Supplementary Methods Figure 4D, we show the model predictions for number of cells and cell apical area depending on the lumen radius. The simulation results are in an excellent agreement with the experimental data and show that the cells have to proliferate rapidly in large epiblasts, whereas the apical area of individual cells remains relatively constant in large epiblasts. This again indicates that in matured epiblasts, apical growth alone cannot drive lumen growth and that there has to be luminal osmotic pressure that overcomes the pressure from the surrounding gel and promotes cell division due to the stretching in the cell layer.

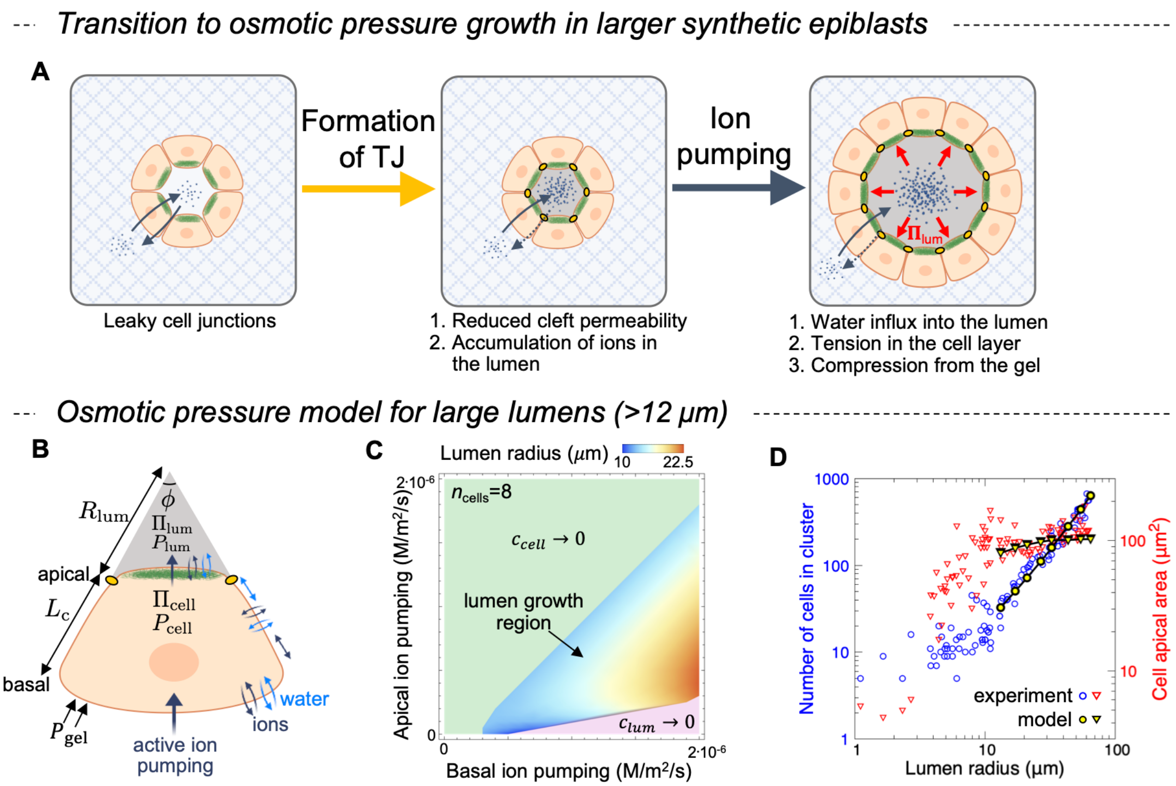

**Supplementary Methods Figure 4: Physical model of osmotic pressure driven lumen growth in large synthetic epiblasts. (A)** Summary of lumen growth mechanism in larger synthetic epiblasts. **(B)** Schematics of the cell and lumen geometry in the model for osmotic pressure driven lumen growth. **(C)** Dependence of lumen size on ion pumping through the basal and apical side of the cell. The region of predicted lumen growth (colored domain) is limited by regions where lumens do not form due lack of ions in the cell (green region) or in the lumen (purple region). **(D)** Model predictions closely match experimental observations of lumen radius, number of cells and cell apical surface area of larger epiblasts.

Physical parameters used in simulations are listed in Supplementary Methods Table 1.

**Supplementary Methods Table 1: Model parameters**

| Parameter | Description | Value |
|---|---|---|
| $R_b^{ini}$ | initial epiblast size [μm] | 12 |
| $L_c^{ini}$ | initial cleft length [μm] | 12 |
| $h_c$ | cleft width [nm] | 20 |
| $P_{ext}$ | external pressure [kPa] | 100 |
| $c_{ext}$ | external ion concentration [mM] | 300 |
| $\omega$ | ion permeability of the cell membrane [mol.m$^{-2}$s$^{-1}$Pa$^{-1}$] | 1.5×10$^{-9}$ |
| $D$ | ion diffusivity in the cleft [m$^2$s$^{-1}$] | 2×10$^{-9}$ |
| $\Lambda_m$ | water permeability of the cell membrane [mol.m$^{-2}$s$^{-1}$Pa$^{-1}$] | 7×10$^{-12}$ |
| $\Lambda_{leak}$ | water permeability of the cleft [mol.m$^{-2}$s$^{-1}$Pa$^{-1}$] | 2×10$^{-14}$ |
| $K$ | effective stiffness of the cortical layer [kPa] | 6 |
| $E_{gel}$ | Young's modulus of the gel [kPa] | 1 |
